# Comparison of different genotyping techniques to distinguish recrudescence from new infection in studies assessing the efficacy of antimalarial drugs against *Plasmodium falciparum*

**DOI:** 10.1101/2023.04.24.538072

**Authors:** Annina Schnoz, Carla Beuret, Maura Concu, Salome Hosch, Liliana K. Rutaihwa, Monica Golumbeanu, Christian Nsanzabana

## Abstract

Distinguishing recrudescence from new infections is crucial for the assessment of antimalarial drug efficacy against *Plasmodium falciparum (P. falciparum)*. Different genotyping methods are used and may impede the comparison of drug efficacy estimates in space and time, particularly in patients from high transmission settings with polyclonal infections.

We compared five different genotyping methods currently used to assess their sensitivity in detecting minority clones in polyclonal infections, their robustness, and the genetic diversity of the markers used. Our study utilized four well-characterized *P. falciparum* laboratory strains mixed in varying ratios, and 20 paired patient samples collected from a clinical trial.

We found that high-resolution capillary electrophoresis (H-CE) using length-polymorphic markers, as well as targeted amplicon deep sequencing (TADS) using single nucleotide polymorphism (SNP)-rich markers, revealed the highest sensitivity in detecting minority clones, while also exhibiting robustness, and high genetic diversity in the used markers. Moreover, markers used by TADS gave more consistent results. We observed that microsatellites had a lower genetic diversity compared to markers such as msp1, *msp2, glurp* and SNP-rich markers, with some genotypes having allelic frequencies of > 30 %.

The replacement of *glurp* by microsatellites did not result in a change in the genotyping outcome, probably due to the lower genetic diversity of microsatellites used in comparison to *glurp*. More studies with large sample sizes are necessary to identify the most suitable microsatellites that could replace *glurp* as per the latest recommendations from the World Health Organization (WHO) on genotyping to distinguish recrudescence from new infections in high transmission settings. Our study indicates that TADS should be considered the gold standard for genotyping to differentiate recrudescence from new infection and should be used to validate other techniques.

## INTRODUCTION

The emergence and spread of partial artemisinin resistance in South east Asia and Africa [1, 2], and subsequent treatment failures observed in South East Asia [3, 4] is threatening the gains made over the last decade in reducing malaria morbidity and mortality to unprecedented low levels [5]. There is a need to strengthen antimalarial drug efficacy surveillance through routine therapeutic efficacy studies to monitor the emergence and spread of artemisinin-based combination therapies (ACTs) resistance.

The assessment of antimalarial drug efficacy requires “PCR correction” to distinguish recrudescence from new infection, especially in high transmission settings where patients are frequently re-infected [6]. Even though the World Health Organization (WHO) has established a standardized protocol for this purpose, various laboratories use different methods with a multitude of molecular markers [7] and different algorithms, leading to difficulties in comparing results over space and time. Initially, the WHO recommended using capillary electrophoresis to genotype three length polymorphic markers: merozoite surface protein 1 (*msp1*), merozoite surface protein 2 (*msp2*), and glutamate-rich protein (*glurp*) [6]. However, other capillary electrophoresis-based assays using other length polymorphic markers such as microsatellites [8], or TaqMan™ Real-time PCR assay using genome-wide single nucleotide polymorphisms (SNPs) barcode [9, 10] have been also used to distinguish recrudescence from new infection. More recently, high resolution melting (HRM) quantitative real-time PCR using *msp1* and *mps2* [11], as well as targeted amplicon deep sequencing (TADS) using single nucleotide polymorphism (SNP)-rich markers [12, 13] have been developed.

All these methods have their advantages, but also limitations that can provide suboptimal results [14]. Some of these limitations, such as the preferential amplification of the smallest amplicons in fragment sizing by capillary electrophoresis have already been explored [15], and this has led the WHO to update its recommendation by replacing the marker *glurp* with microsatellites [16]. Even so, the optimal microsatellite to replace *glurp* is yet to be described and characterized. Recently developed methods such as TADS have the potential to overcome some of the limitations of current methods and improve the accuracy and robustness of the genotyping outcomes. However, all these methods have varying technical requirements and running costs [17], and there has been no systematic head-to-head comparison of these assays to assess their sensitivity, robustness, genetic marker diversity and impact on genotyping outcomes with the aim of improving comparability between laboratories. In this study, we compared five methods to distinguish recrudescence from new infection using the same samples and the same operator in a single laboratory.

## METHODS

### Laboratory cultured parasite strains

*Plasmodium falciparum* laboratory strains (3D7, K1, HB3 and FCB1) were obtained from BEI Resources and cultured according to standard protocol [18]. Parasites were harvested at 5% parasitemia after having been synchronized by sorbitol to obtain ring stages.

### Patient samples

Samples were collected during a clinical trial to evaluate the efficacy and safety of Cipargamin (KAE609) in a randomized, Phase II dose-escalation study in adults in with uncomplicated *P. falciparum* malaria. The trial was conducted in high transmission settings in sub-Saharan Africa. Details about the study protocol and patients’ recruitment have been published elsewhere [19]. Twenty paired samples from patients presenting with parasite recurrence during the clinical trial were selected for the study.

### DNA extraction

DNA was extracted from *P. falciparum* parasite culture and from malaria patients using the QIAsymphony DSP DNA Mini Kit according to the manufacturer instructions [20].

### Preparation of laboratory strains mixes

Parasitaemia of the synchronized ring-stage parasite strains 3D7, K1, HB3 and FCB1 was measured by qPCR targeting 18s rRNA gene as previously described [21]. The four laboratory strains were mixed in different ratios and diluted in human DNA of malaria negative donors (Supplementary Table 1). DNA from the four strains was mixed in different ratios ranging from 1:1:1:1 to 3000:1:1:1, and the concentration of the minority clone was always 10 parasites/μl as previously described [15]. In total, 38 different strain mixtures were prepared (Supplementary Table 1) and each mixture was tested in three technical triplicates. The analysis was repeated by the same operator on a different day to assess the robustness of the different assays in detecting minority clones.

### Msp1, msp2 and glurp genotyping using fast (F-CE) – and high-resolution capillary electrophoresis (H-CE)

*Msp1, msp2* and *glurp* were amplified by primary and nested PCR as previously described [15]. The sizes of nested PCR products were assessed by F-CE using Qiaxcel®, and H-CE using ABI 3730xl®. A 10% cut-off for *msp1* and *msp2* and a 20% cut-off for *glurp* were applied to remove stutter peaks as previously described [15]. Detailed protocols are described in Supplementary Methods 1 and 2.

### Microsatellites genotyping using high-resolution capillary electrophoresis (H-CE)

Four different microsatellites (PfPK2, TA40, TA60 and TA81) were amplified in a single-round PCR as previously described [8]. The size of the PCR products was assessed by H-CE using ABI 3730xl® and peaks were analyzed using a microsatellites-specific cut-off as previously described [8]. Detailed protocols are provided in Supplementary Methods 3.

### SNPs-rich markers genotyping using targeted amplicon deep sequencing (TADS)

Five different markers, *ama1-D3, cpmp, cpp, csp* and *msp7*, were amplified by primary and nested PCR, followed by adapter PCR, magnetic purification and normalization as previously described [12, 13]. Sequencing was done on a Miseq in paired-end mode using the MiSeq reagent kit v3 (600 cycles), and run with 15% spike-in of Enterobacteria phage *phiX* control v3 [12]. Detailed protocols are described in Supplementary Methods 4.

### Msp1 and msp2 genotyping using high-resolution melting (HRM)

Quantitative PCR was performed followed by DNA melting in simplex assays with specific primers for *msp1* and *msp2*, as previously described [11]. Detailed protocol about PCR conditions and data analysis can be found in Supplementary Methods 5.

### Data analysis

The limit of detection (LOD) for minority clones was defined as the ratio of laboratory strains returning two out of three replicates positive. To assess the intra assay variability, we calculated the percentage of samples that returned the three technical replicates with the same result, and the inter assay variability as the percentage of pairs of technical replicates giving the same LOD between two runs. The genetic diversity was assessed by calculating the number of genotypes (number of fragments in electropherogram of each sample for length polymorphic markers; number of haplotypes found per sample in TADS), and their frequency in the 40 samples (20 pairs) collected from the clinical trial. The scripts used for automated microsatellite cut-off analysis, TADS SNPs – and haplotype calling as well as pie charts and bar plots can be found under the following link: https://github.com/SwissTPH/AmpSeqAnalysis. Two different algorithms were used to interpret the genotyping results with several markers. The WHO algorithm defines as recrudescence, sample pairs showing a recrudescence for at least one marker [6]. The two out of three algorithm defines as recrudescence, only sample pairs for which the majority of markers used are showing a recrudescence [15].

## RESULTS

### Limit of detection for minority clones and reproducibility

#### *Msp1, msp2* and *glurp* genotyping using fast (F-CE) – and high-resolution capillary electrophoresis (H-CE)

The limit of detection (LOD) for minority clones varied between the different *msp1* and *msp2* allelic families as well as between assays when the minority clone had the longest amplicon (Figure 1). As the four strains used fall in two allelic families, the comparison for *msp1* and *msp2* allelic families could be only done for two strains. Overall, compared to F-CE, higher sensitivity was observed with H-CE for almost all markers. The LOD increased 20 times for 3D7 and 6 times for Mad20 when using H-CE compared to F-CE, going from detecting minority clones in 1 in 1 ratio to 1 in 20 ratio, and from 1 in 15 to 1 in 100 for 3D7 and Mad20 respectively (Figure 1). The LOD was the highest with Mad20 allelic family for both F-CE and H-CE and was the same for K1 allelic family with both assays (Figure 1). For *glurp*, the longest amplicons could not be detected when they were in minority. The intra- and inter-assay variability were higher for F-CE compared to H-CE (Figure 2). The reproducibility varied between the different markers for F-CE, especially the inter-assay reproducibility being as low as 37% for K1 but was consistently high and close to 100% for H-CE (Supplementary Table 2 and 3). The reproducibility for *glurp* was high for both F-CE and H-CE due to the fact that minority clones were never detected.

**Figure 1.**
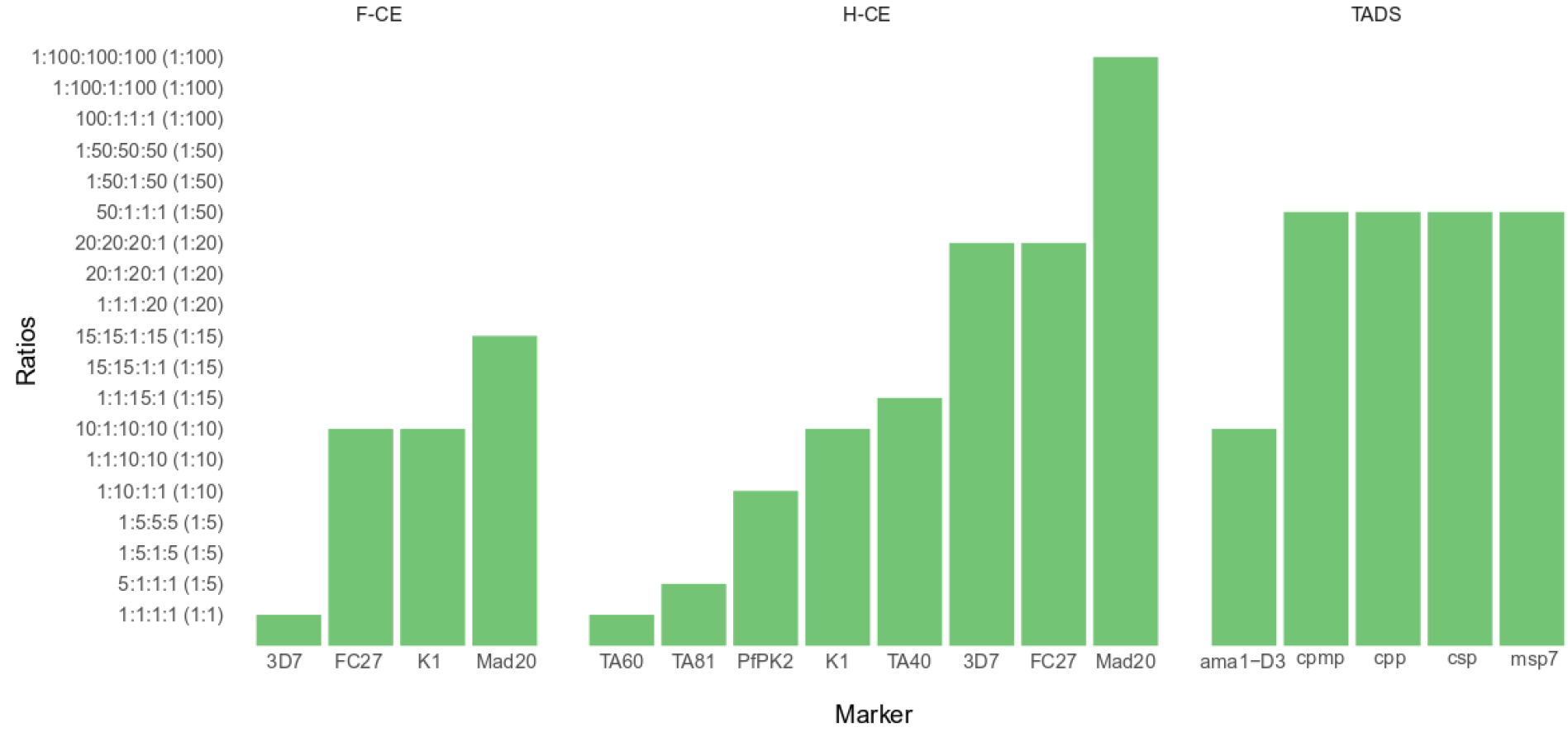
Limit of detection for minority clones for different genotyping techniques and different markers used to differentiate recrudescence from new infection. Fast capillary electrophoresis (F-CE) using *msp1* and *msp2* with long amplicon in minority, high-resolution capillary electrophoresis (H-CE) using *msp1* and *msp2* and microsatellites with long amplicon in minority, targeted amplicon deep sequencing (TADS) using SNP-rich markers. Y-axis: *P. falciparum* laboratory strains mix ratios [3D7:K1:HB3:FCB1)]

**Figure 2.**
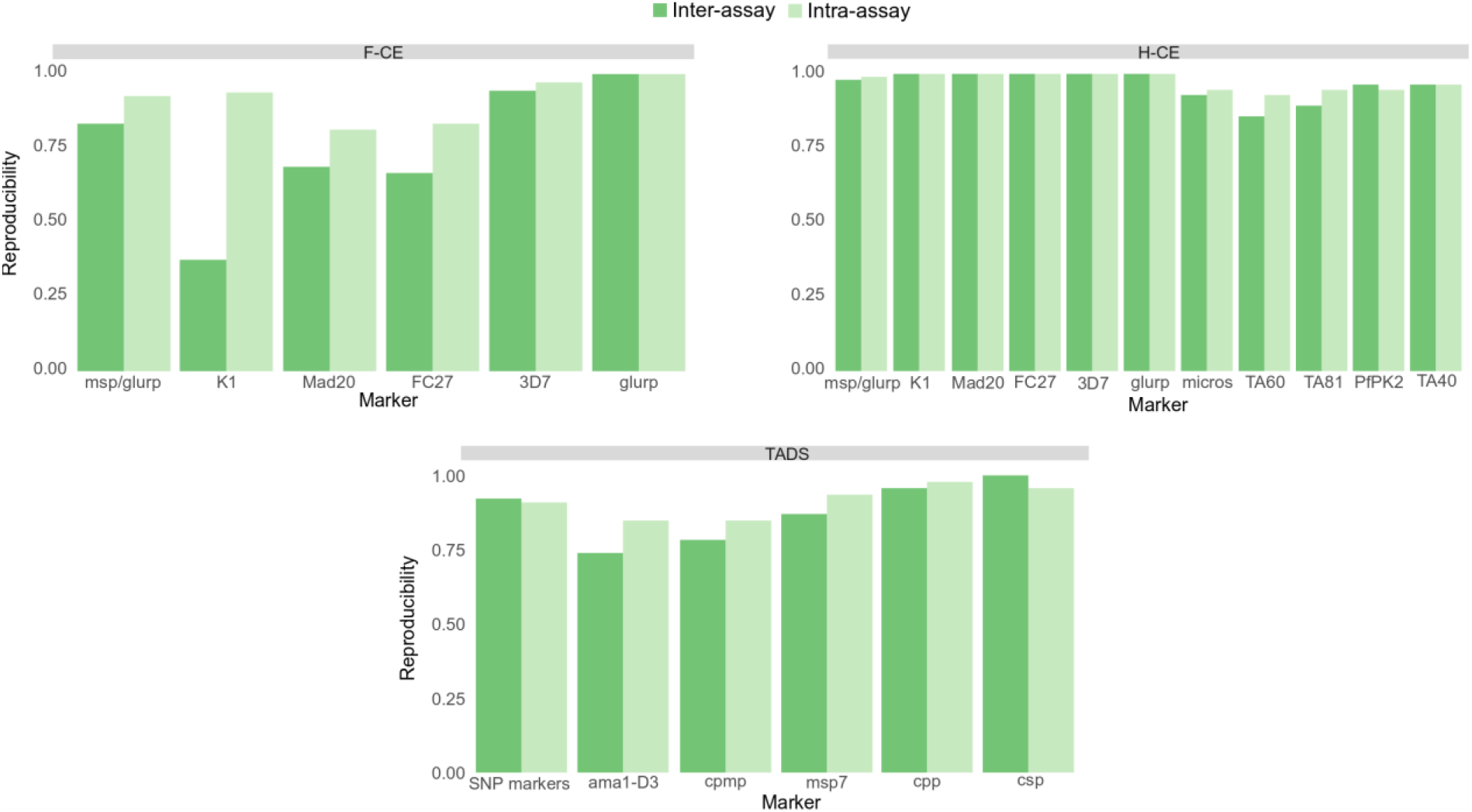
Inter- and intra-assay reproducibility. Inter-assay reproducibility: percentage of pairs of technical replicates are concordant on the limit of detection. Intra-assay reproducibility: percentage of samples for which the three technical replicates are concordant on the limit of detection

#### Microsatellites *(PfPk2, TA40, TA60, TA81)* genotyping using high-resolution capillary electrophoresis (H-CE)

Analysis of microsatellites was only performed with H-CE due to the higher resolution, as microsatellites may differ only in 3 base pairs. The LOD was lowest for TA60 and TA81 (1 in 1 and 1 in 5 ratios, respectively) and the highest for TA40 (1 in 15 ratio) (Figure 1). The limit of detection for *PfPk2* (1 in 10 ratio) was similar to the LOD for K1 (*msp2*). However, for the different microsatellites, there were always two laboratory strains with the same amplicon size, except for TA81 (Supplementary Table 4). The reproducibility for microsatellites was consistently above 90%, except the inter-assay reproducibly for TA60 that was 85.7% (Figure 2).

#### SNPs-rich markers *(ama1-D3, cpmp, cpp, csp, msp7)* genotyping using targeted amplicon deep sequencing (TADS)

The LOD was identical for 4 markers, except for *ama1-D3*, which had an LOD approximately half of that of the other markers (Figure 1). *Cpmp, cpp, csp*, and *msp7* demonstrated the ability to detect minority clone when the majority clone was present at a concentration 20 times higher, similar to the performance of H-CE with 3D7 and FC27. On the other hand, *ama1-D3* was able to detect a minority clone with a concentration 10 times lower than that of the majority clone, which is comparable to the performance of K1. Both *cpp* and *csp* showed very high levels of intra- and inter-assay reproducibility, exceeding 95%. However, reproducibility was lower for *msp7, cpmp*, and *ama1-* D3 being below 90%. (Figure 2).

#### *Msp1 and msp2* genotyping using high-resolution melting analysis (HRM)

In both runs and for both markers, none of the samples showed more than two different melting temperatures within the expected range. Depending on the marker, two to three strains had almost identical melting temperatures for each marker resulting in overlap of mixed-strain sample peaks.

### Markers diversity and allelic frequency

The diversity, defined as the number of distinct genotypes, exhibited variation across different markers (Figure 3). Among the markers, four had more than 30 distinct genotypes (3D7 allelic family, FC27 allelic family, *glurp*, and *cpmp*), with the 3D7 allelic family having the highest number of genotypes (36). Additionally, five markers had more than 20 genotypes (K1 allelic family, Mad20 allelic family, *ama1-D3, cpp* and *csp*), with K1 and *ama1-D3* having the highest number of genotypes (28). The microsatellites markers TA60 and TA81 showed the lowest number of genotypes, with both markers harboring only 9 genotypes. The allelic frequency of the genotypes varied across the different markers. Two markers (*cpmp* and FC27) did not have any genotype with an allelic frequency of ≥10%. The 3D7 and K1 allelic families had only one genotype with allelic frequency of 10%, whereas TA60 and *msp7* had genotypes with allelic frequencies >30%. All microsatellites had at least one genotype with an allelic frequency of >20%.

**Figure 3.**
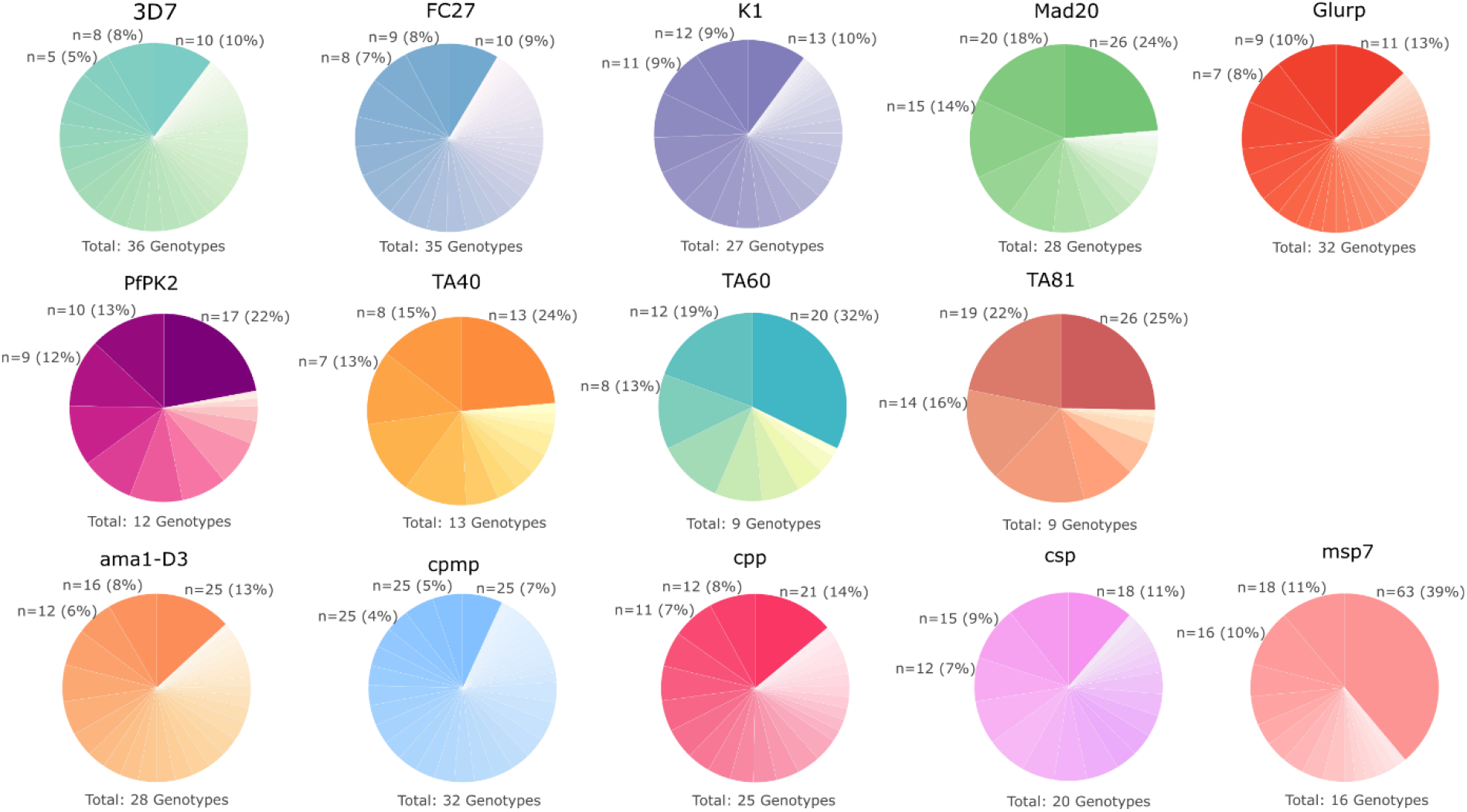
Diversity of the different length polymorphic markers (*msp2* allelic families: 3D7 and FC27, *msp1* allelic families: K1 and Mad20, *glurp*, and microsatellites: *PfPk2, TA40, TA60, TA81*) assessed with high-resolution capillary electrophoresis (H-CE) and single nucleotide polymorphisms (SNPs-rich markers (*ama1-D3, cpmp, cpp, csp, msp7*) assessed with targeted amplicon deep sequencing (TADS)

### Distinguishing recrudescence (R) from new Infection (NI) in an antimalarial drug clinical trial

#### Concordance between markers

Twenty patients with recurrent infections enrolled in a clinical trial for the drug Cipargamin were included in this study, and their samples were genotyped using four different assays. All techniques and markers yielded concordant results for 10 patients (Table 1). When each assay was evaluated independently, TADS showed higher concordance between markers, with all markers giving the same outcome for 90% of the patients. The lowest concordance between markers was observed with F-CE, with only 65% of patients showing the same outcome for all markers. All baseline and recurrent samples were genotyped for the *PfATP4* G538S mutation that confers clinically relevant resistance to Cipargamin. Of the 16 patients that a had a recrudescence confirmed by TADS, 14 of them (87.5%) had the *PfATP4* G538S mutation in the recrudescent samples, but not in the baseline samples collected before treatment. None of the four patients with a new infection confirmed by TADS, had this mutation.

**Table 1.**
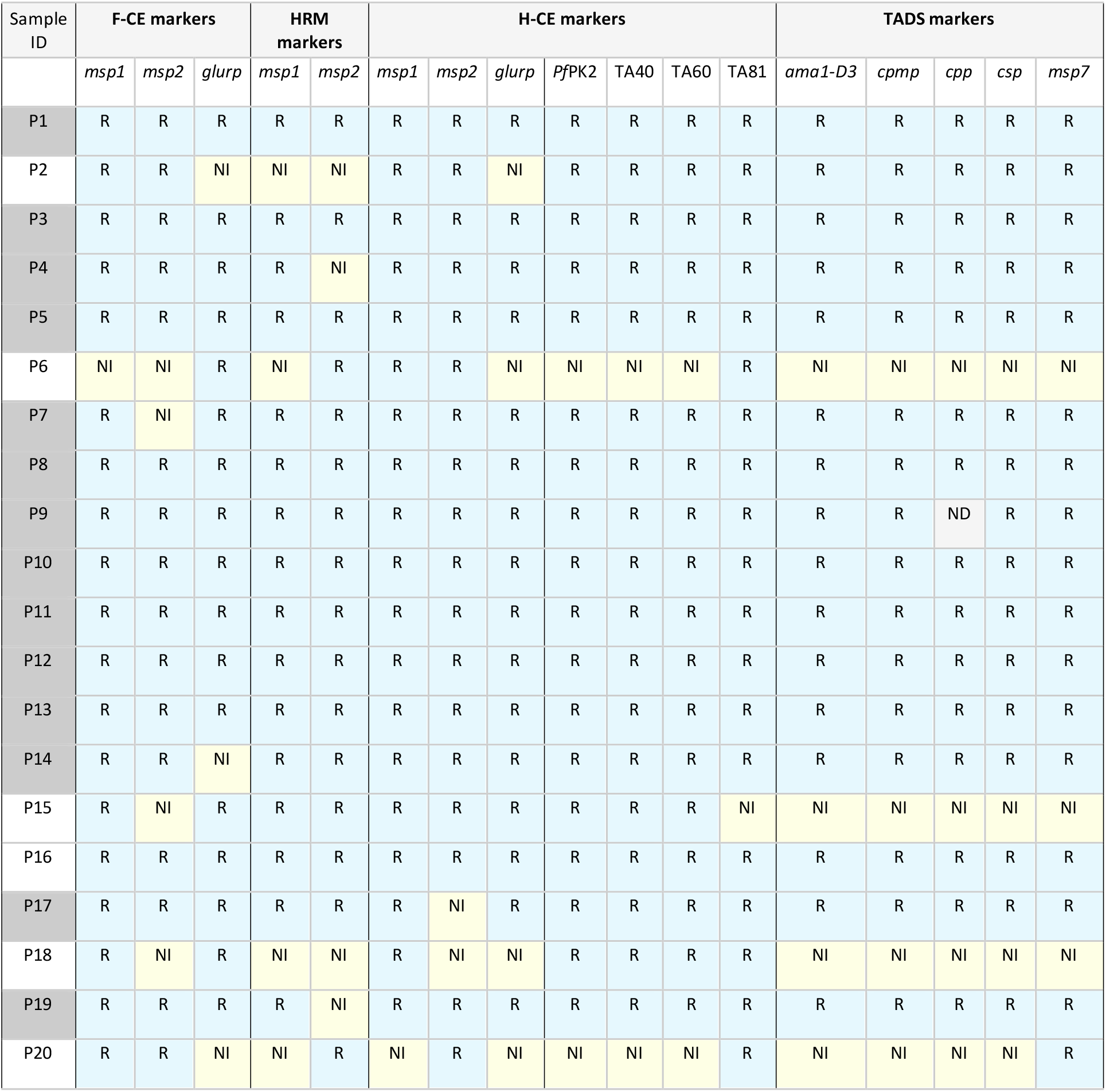
New infection and recrudescence outcome of the different methods and their markers using 20 paired patient samples. R = recrudescence outcome (blues shading); NI = new infection outcome (yellow shading); ND = not determined. Patients shaded in grey: recurrent samples with the *PfATP4* G538S mutation associated with resistance to Cipargamin.

#### Concordance between results interpretation algorithms

For the 10 patients with discordant results between the different markers and different assays, the final genotyping outcome (recrudescence or new infection) was determined using two different algorithms: the WHO algorithm and 2 out of 3 algorithm (Table 2). The results showed that for TADS, the outcome of both algorithms was concordant for all 10 patients. Six patients had recrudescent infections by TADS for both algorithms. However, for H-CE using *msp1, msp2* and *glurp* as well as *msp1, msp2* and microsatellites, the two algorithms agreed on the outcome in 7 out of 10 patients. From the three discordant cases, two were due to the marker *glurp*. When using the 2 out of 3 algorithm, the number of recrudescent infections increased slightly for H-CE with *msp1/msp2/glurp* or *msp1/msp2*/microsatellites comparing to the WHO algorithm (Table 2). In contrast, F-CE and HRM found a substantial lower number of recrudescent infections, with only three and four, respectively with the WHO algorithm. However, when using the 2 out of 3 algorithm, the number of recrudescent infections increased to nine for H-CE F-CE. It was not possible to use the 2 out 3 algorithm for *msp1/msp2* HRM, as only two markers were used.

**Table 2.** Final outcome using **WHO** and **2/3** algorithm (also 4/5 for TADS and 4/6 algorithm for H-CE *msp1, msp2* and four microsatellites) of ten sample pairs with discrepant results between markers. For the HRM method no 2/3 algorithm was applied because only two markers were used.

| Sample ID | F-CE<br>(*msp1*, *msp2*,<br>*glurp*) |  | HRM<br>(*msp1*,<br>*msp2*) | H-CE<br>(*msp1*, *msp2*,<br>*glurp*) |  | H-CE<br>(*msp1*, *msp2*,<br>PfPk2 or TA40 or<br>TA60) |  | H-CE<br>(*msp1*, *msp2*,<br>TA81) |  | TADS<br>(*ama1-D3*, *cscp*,<br>*cpp*, *cpmp*, *msp7*) |  |
| --- | --- | --- | --- | --- | --- | --- | --- | --- | --- | --- | --- |
|  | WHO | 2/3 | WHO | WHO | 2/3 | WHO | 2/3 | WHO | 2/3 | WHO | 2/3 |
| S2 | NI | R | NI | NI | R | R | R | R | R | R | R |
| S4 | R | R | NI | R | R | R | R | R | R | R | R |
| S6 | NI | NI | NI | NI | R | NI | R | R | R | NI | NI |
| S7 | NI | R | R | R | R | R | R | R | R | R | R |
| S14 | NI | R | R | R | R | R | R | R | R | R | R |
| S15 | NI | R | R | R | R | R | R | NI | R | NI | NI |
| S17 | R | R | R | NI | R | NI | R | NI | R | R | R |
| S18 | NI | R | NI | NI | NI | NI | R | NI | R | NI | NI |
| S19 | R | R | NI | R | R | R | R | R | R | R | R |
| S20 | NI | R | NI | NI | NI | NI | NI | NI | R | NI | NI |
| Total number of<br>recrudescences | 3 | 9 | 4 | 5 | 8 | 6 | 9 | 7 | 8 | 6 | 6 |

## DISCUSSION

PCR correction is essential for accurately estimating antimalarial drug efficacy, especially in high transmission settings. However, different assays and different molecular markers are used to differentiate recrudescence from new infection, making it difficult to compare results from different laboratories. The optimal assay should be highly sensitive in detecting minority clones in polyclonal infections, have a high intra- and inter-assay reproducibility, and use molecular markers with a high diversity and low allelic frequencies of the different genotypes [22].

To our knowledge, this is the first study to systematically compare the most used assays to distinguish recrudescence from new infection [7]. We assessed their ability to detect minority clones in polyclonal infections, their robustness, the genetic diversity and allelic frequency of the markers used. As previously described, SNPs-rich markers analyzed by TADS, and *msp1*/*msp2* analyzed by H-CE proved to be the most sensitive assays in detecting minority clones in polyclonal infections [12]. Moreover, these two techniques are very robust, with high intra and inter-assay reproducibility. The lower reproducibility of some TADS markers emphasizes the need for high sequencing coverage and analysis of samples in triplicates to get consistent results as previously described [13]. The markers used by these two techniques also have the highest diversity, low allelic frequencies, with few genotypes having a frequency over 10% in samples collected from high transmission settings. However, TADS has an additional advantage, the five markers used were consistently giving concordant results in most of samples analyzed, making the final outcome not being dependent on the algorithm used to analyze the data. This is in contrast to H-CE, for which the discordance between markers sometimes led to different final outcomes, when different algorithms were used for data analysis. Previous studies have also shown that TADS has higher concordance between markers than capillary electrophoresis [12]. The reason behind this could be the presence of stutter peaks in H-CE, which are challenging to differentiate from real amplicons, and may result to false genotyping [23].

The most recent WHO guidelines for distinguishing recrudescence from new infection suggest using the best microsatellites in combination with *msp1* and *msp2* to replace *glurp* [16], due to the low sensitivity of *glurp* in detecting minority clones [15], which was also confirmed in our study. Here, we assessed four different microsatellites (*PfPk2, TA40, TA60*, and *TA81*), that have were previously demonstrated high sensitivity in detecting minority clones and high genetic diversity [8]. All of these markers were found to have higher sensitivity than *glurp* in detecting minority clones, but their genetic diversity was lower. In the 40 samples analyzed, there were 32 different genotypes for *glurp*, but only nine to 13 for microsatellites. Moreover, some genotypes in microsatellites had high allelic frequencies up to 32%, and only one genotype in *glurp* had an allelic frequency > 10%. Due to its low sensitivity in detecting minority clones, it is expected that *glurp* may fail to detect recrudescent infections at low levels, leading to underestimation of the true number of recrudescences. Conversely, the lower genetic diversity and high allelic frequency of certain genotypes in some microsatellites may lead to an overestimation of recrudescent infections. This is in line with our findings, as in the 20 sample pairs analyzed, *glurp* identified four new infections, and two of them were identified as recrudescences by microsatellites. However, due to the limited sample size, further assessment of other microsatellites is necessary to draw a definitive conclusion.

The WHO currently recommends a sequential algorithm, which gives the final result as a new infection when one of the markers detects a new infection [6, 16]. However, this algorithm may not be optimal, particularly when the sensitivity of the markers in detecting minority clones differs. This could lead to an underestimation of recrudescences. Therefore a 2 out of 3 algorithm which takes into account the two markers that are in agreement, has been suggested to improve accuracy [24]. In our study, we used TADS as the reference method for determining recrudescence, due to its high level of robustness and concordance between markers, and its demonstrated association with “true recrudescence”. To identify a “true recrudescence” can be challenging, and therefore surrogate markers need to be utilized, such as molecular markers associated with antimalarial drug resistance [8]. Here, we utilized the *PfATP4* G538S mutation as a molecular marker, which has been previously associated with resistance to Cipargamin both *in vitro* and *in vivo* [19, 25]. In a phase II clinical trial, 65% of recrudescent infections were found to have this specific mutation, which was not present in samples collected before treatment [19]. In our sample set, 14 out of 16 recrudescent infections were found to have this mutation. On the other hand, none of the new infections determined by TADS had this mutation. These results give us high confidence in the genotyping results obtained from this method. We evaluated the performance of various genotyping assays and markers using two different algorithms (WHO and two out of three) on a set of 20 sample pairs. Although the sample size was small, our findings suggest that replacing *glurp* with microsatellites may not lead to a significant improvement in the final genotyping outcome. The low sensitivity of *glurp* in detecting minority clones could be compensated for by its high diversity and low allelic frequencies. The significant discrepancy between the two algorithms for F-CE (3 and 9 recrudescences for WHO and 2 out of 3 algorithms, respectively) indicates a lack of discriminatory power of this technique, similar to agarose gels. This highlights the importance of using techniques with high discriminatory power such as H-CE and TADS. Previous studies have also reported low discriminatory power of agarose gels compared to H-CE in samples from high transmission settings with high multiplicity of infection [26, 27], which is consistent with our findings. Algorithms based on Bayesian statistics have been developed for microsatellites and have the potential to improve the accuracy of the final outcome by taking into account the different parameters assessed in this study such as sensitivity in detecting minority clones, robustness, markers genetic diversity and allelic frequency for each technique [28]. They provide uncertainty estimates and give a different weight to each marker based on its performance, potentially providing a more accurate final result [29].

There are some limitations to this study that should be acknowledged. Firstly, we only used technical replicates extracted from parasite culture or patients’ samples once, which may not be sufficient to fully capture the stochastic variations that could arise from DNA extraction. Secondly, the LOD for *msp1* and *msp2* was only assessed using two strains, as two of the laboratory strains belonged to the same allelic family. Similarly, for microsatellites, there were always two strains with the same amplicon length, except for *TA81* where the four strains possessed four different amplicon sizes. Finally, the samples set used to differentiate between recrudescence and new infection was small and had a high proportion of recrudescence, limiting the generalizability of our results.

In summary, our study suggests the recent changes in WHO guidelines for genotyping may not lead to improved outcomes due to the lower genetic diversity of microsatellites compared to *glurp*. Further studies with larger sample sizes and other microsatellites are needed to identify the best replacements for *glurp*. Moreover, high-resolution techniques such as H-CE should be preferred for length polymorphic markers instead of techniques with lower resolutions such as F-CE or agarose gels. TADS should be considered as the gold standard in the future for validating new techniques due to its robustness, and high consistency in results across markers.

## Acknowledgements

We would like to thank Dr. Khalid Beshir from LSHTM to have shared his protocol for *msp1, msp2* HRM and support for implementing the assay in our laboratory. We also thank Armin Passecker from Swiss TPH for his support for laboratory strains culture and Maria Gruenberg for her support in establishing the amplicon deep sequencing assay. Our thanks go also to Prof. Pascal Mäser who co-supervised Annina Schnoz during her Msc thesis, and to Dr. Anita Lerch for her support in data analysis with HaplotypR. We would like to extend our acknowledgements to Dr. Esther Schmitt from Novartis Pharma AG, Basel for providing samples that were used in this study, and Prof. Ingrid Felger for her critical reading of the manuscript.

