## Supplemental file 1 for "Comparison of different genotyping techniques to distinguish recrudescence from new infection in studies assessing the efficacy of antimalarial drugs against *Plasmodium falciparum*"

### SUPPLEMENTARY FILES

**Supplementary Table 1.** Lab-strain mixtures with different ratios used to assess the genotyping methods.

| SAMPLE ID | STRAIN RATIO | | | | PARASITES PER µL | | | |
|---|---|---|---|---|---|---|---|---|
| | 3D7 | K1 | HB3 | FCB1 | 3D7 | K1 | HB3 | FCB1 |
| **S 1** | 1 | 0 | 0 | 0 | 10 | 0 | 0 | 0 |
| **S 2** | 0 | 1 | 0 | 0 | 0 | 10 | 0 | 0 |
| **S 3** | 0 | 0 | 1 | 0 | 0 | 0 | 10 | 0 |
| **S 4** | 0 | 0 | 0 | 1 | 0 | 0 | 0 | 10 |
| **S 5** | 1 | 1 | 1 | 1 | 10 | 10 | 10 | 10 |
| **S 1 NEW** | 1 | 0 | 0 | 0 | 1'000 | 0 | 0 | 0 |
| **S 2 NEW** | 0 | 1 | 0 | 0 | 0 | 1'000 | 0 | 0 |
| **S 3 NEW** | 0 | 0 | 1 | 0 | 0 | 0 | 1'000 | 0 |
| **S 4 NEW** | 0 | 0 | 0 | 1 | 0 | 0 | 0 | 1'000 |
| **S 5 NEW** | 1 | 1 | 1 | 1 | 1'000 | 1'000 | 1'000 | 1'000 |
| **S 6** | 5 | 1 | 1 | 1 | 50 | 10 | 10 | 10 |
| **S 7** | 1 | 5 | 1 | 5 | 10 | 50 | 10 | 50 |
| **S 8** | 1 | 5 | 5 | 5 | 10 | 50 | 50 | 50 |
| **S 9** | 1 | 10 | 1 | 1 | 10 | 100 | 10 | 10 |
| **S 10** | 1 | 1 | 10 | 10 | 10 | 10 | 100 | 100 |
| **S 11** | 10 | 1 | 10 | 10 | 100 | 10 | 100 | 100 |
| **S 12** | 1 | 1 | 15 | 1 | 10 | 10 | 150 | 10 |
| **S 13** | 15 | 15 | 1 | 1 | 150 | 150 | 10 | 10 |
| **S 14** | 15 | 15 | 1 | 15 | 150 | 150 | 10 | 150 |
| **S 15** | 1 | 1 | 1 | 20 | 10 | 10 | 10 | 200 |
| **S 16** | 20 | 1 | 20 | 1 | 200 | 10 | 200 | 10 |

| | | | | | | | | |
|---|---|---|---|---|---|---|---|---|
| **S 17** | 20 | 20 | 20 | 1 | 200 | 200 | 200 | 10 |
| **S 18** | 50 | 1 | 1 | 1 | 500 | 10 | 10 | 10 |
| **S 19** | 1 | 50 | 1 | 50 | 10 | 500 | 10 | 500 |
| **S 20** | 1 | 50 | 50 | 50 | 10 | 500 | 500 | 500 |
| **S 21** | 100 | 1 | 1 | 1 | 1'000 | 10 | 10 | 10 |
| **S 22** | 1 | 100 | 1 | 100 | 10 | 1'000 | 10 | 1000 |
| **S 23** | 1 | 100 | 100 | 100 | 10 | 1'000 | 1'000 | 1'000 |
| **S 24** | 1 | 500 | 1 | 1 | 10 | 5'000 | 10 | 10 |
| **S 25** | 1 | 1 | 500 | 500 | 10 | 10 | 5'000 | 5'000 |
| **S 26** | 500 | 1 | 500 | 500 | 5'000 | 10 | 5'000 | 5'000 |
| **S 27** | 1 | 1 | 1000 | 1 | 10 | 10 | 10'000 | 10 |
| **S 28** | 1000 | 1000 | 1 | 1 | 10'000 | 10'000 | 10 | 10 |
| **S 29** | 1000 | 1000 | 1 | 1000 | 10'000 | 10'000 | 10 | 10'000 |
| **S 30** | 1 | 1 | 1 | 1500 | 10 | 10 | 10 | 15'000 |
| **S 31** | 1500 | 1 | 1500 | 1 | 15'000 | 10 | 15'000 | 10 |
| **S 32** | 1500 | 1500 | 1500 | 1 | 15'000 | 15'000 | 15'000 | 10 |
| **S 33** | 3000 | 1 | 1 | 1 | 30'000 | 10 | 10 | 10 |
| **S 34** | 1 | 3000 | 1 | 1 | 10 | 30'000 | 10 | 10 |
| **S 35** | 1 | 1 | 3000 | 1 | 10 | 10 | 30'000 | 10 |
| **S 36** | 1 | 1 | 1 | 3000 | 10 | 10 | 10 | 30'000 |
| **S EXTRA 1** | 1 | 1 | 100 | 1 | 10 | 10 | 1'000 | 10 |
| **S EXTRA 2** | 1 | 1500 | 1 | 1 | 10 | 15'000 | 10 | 10 |

3

**Supplementary Table 2.** Division of strains into different allelic families and corresponding fragment sizes.

| Strain | *Msp1* allelic families | *Msp2* allelic families | *Glurp* |
|---|---|---|---|
| **3D7** | K1-type (247 bp) | 3D7-type (264 bp) | glurp-type (881 bp) |
| **K1** | K1-type (177 bp) | FC27-type (407 bp) | glurp-type (822 bp) |
| **HB3** | Mad20-type (157 bp) | FC27-type (335 bp) | glurp-type (537 bp) |
| **FCB1** | Mad20-type (192 bp) | 3D7-type (341 bp) | glurp-type (711 bp) |

**Supplementary Table 3.** Detection limit and robustness of both runs of fast capillary electrophoresis using *msp1* and *msp2* with **long fragment in minority**. For each marker the amplicon size of each strain is indicated in bp.

**LIMIT OF DETECTION FOR DIFFERENT ALLELIC FAMILIES**

| | **K1** (3D7**(247)**:K1**(177)**) | **Mad20** (HB3**(157)**:FCB1**(192)**) | **FC27** (K1**(407)**:HB3**(335)**) | **3D7** (3D7**(264)**:FCB1**(341)**) |
|---|---|---|---|---|
| Run 1 | 1:10 | 20:1 | 1:10 | 1:1 |
| Run 2 | 1:10 | 1:1 | 1:1 | 1:1 |

**Supplementary Table 4.** Detection limit and robustness of both runs of high-resolution capillary electrophoresis using *msp1* and *msp2* with **long fragment minority**, applying the 10% cut-off. For each marker the amplicon size of each strain is indicated in bp.

**LIMIT OF DETECTION FOR DIFFERENT ALLELIC FAMILIES**

| | **K1** (3D7**(247)**:K1**(177)**) | **Mad20** (HB3**(157)**:FCB1**(192)**) | **FC27** (K1**(407)**:HB3**(335)**) | **3D7** (3D7**(264)**:FCB1**(341)**) |
|---|---|---|---|---|
| Runs identical | 1:10 | 100:1 | 1:20 | 20:1 |

**Supplementary Table 5.** Detection limit and robustness of both runs of high-resolution capillary electrophoresis using microsatellite markers with **long fragment in minority** applying the microsatellite cut-offs of Greenhouse et al., 2006. For each marker the amplicon size of each strain is indicated in bp (3D7:K1:HB3:FCB1).

**LIMIT OF DETECTION FOR MICROSATELLITES**

| | ***Pf*PK2** (**247**:**247**:270:241) | **TA40** (270:**249**:255:**249**) | **TA60** (325:**322**:319:**322**) | **TA81** (182:188:191:179) |
|---|---|---|---|---|
| Runs identical | 1:10:1:1 | 1:1:15:1 | 1:1:1:1 | 5:1:1:1 |

**Supplementary Table 6.** Detection limit and robustness of both runs of targeted amplicon deep sequencing using SNP-rich markers. SIM: Strains in minority. For each marker the amplicon size of each strain is indicated in bp (3D7:K1:HB3:FCB1).

**LIMIT OF DETECTION FOR SNP-RICH MARKERS**

| | *ama1D3* | *cpmp* | *cpp* | *csp* | *msp7* |
|---|---|---|---|---|---|
| Run 1: 1/4 SIM | 50:1:1:1 | 50:1:1:1 | 50:1:1:1 | 50:1:1:1 | 1:10:1:1 |
| Run 2: 1/4 SIM | 1:10:1:1 | 100:1:1:1 | 50:1:1:1 | 50:1:1:1 | 50:1:1:1 |
| Run 1: 2/4 SIM | 1:1:10:10 | 1:1:10:10 | 1:1:10:10 | 1:1:10:10 | 1:1:10:10 |
| Run 2: 2/4 SIM | 1:1:10:10 | 1:1:10:10 | 1:1:10:10 | 1:1:10:10 | 1:1:10:10 |
| Run 1: 3/4 SIM | 1:5:5:5 | 10:1:10:10 | 10:1:10:10 | 10:1:10:10 | 1:50:50:50 |
| Run 2: 3/4 SIM | 1:5:5:5 | 10:1:10:10 | 10:1:10:10 | 10:1:10:10 | 10:1:10:10 |

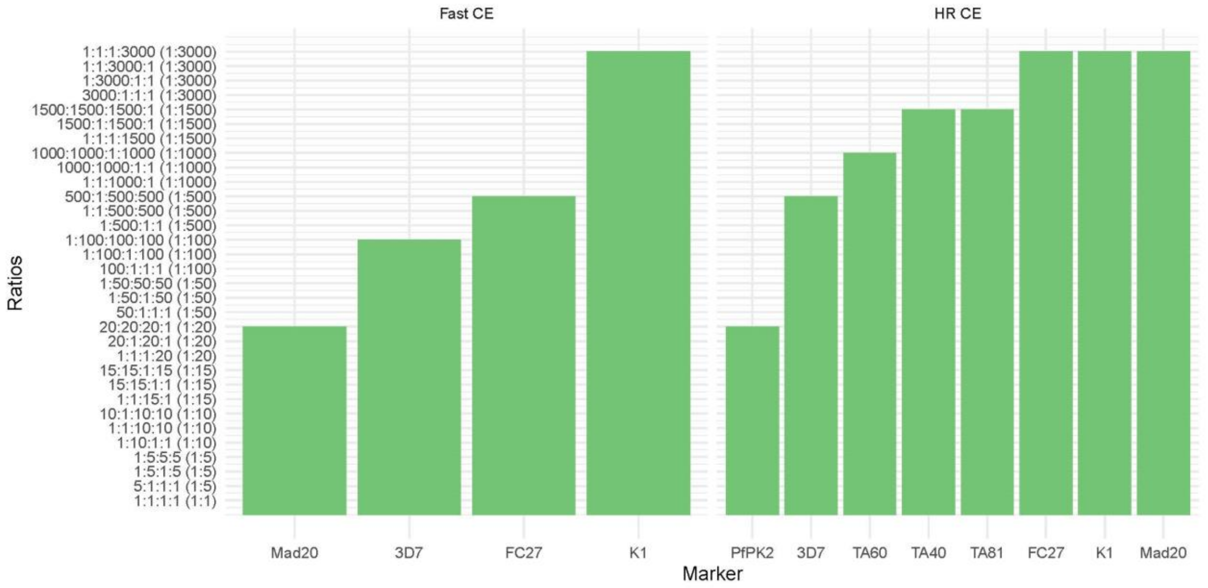

**Supplementary Figure 1.** Detection limit of both runs combined of: fast capillary electrophoresis (F-CE) using *msp1* and *msp2* with **short fragment in minority,** high-resolution capillary electrophoresis (H-CE) using *msp1* and *msp2* (with 10% cut-off) as well as microsatellites (with Greenhouse cut-off) with **short fragment in minority.** Y-axis: strain ratios [3D7:K1:HB3:FCB1)], in base pairs for each marker: **K1** [247:177]; **Mad20** [192:157]; **FC27** [407:335]; **3D7** [341:264]; **TA60** [325:322:319:322]; **TA81** [182:188:191:179]; **TA40** [270:249:255:249]; ***Pf*PK2** [247:247:270:241]. X-axis: allelic family marker.

For *glurp*, the sample harboring **15:15:1:1** ratio [3D7 (**881** bp):K1 (**822** bp):HB3 (**537** bp):FCB1 (**711** bp)] was the only sample positive for all strains, when the 20% cut-off was applied (not shown).

**Supplementary Table 7.** Detection limit and robustness of both runs of fast capillary electrophoresis using *msp1* and *msp2* with **small fragment minority**. For each marker the amplicon size of each strain is indicated in bp.

**LIMIT OF DETECTION FOR DIFFERENT ALLELIC FAMILIES**

| | **K1**<br>(3D7**(247)**:K1**(177)**) | **Mad20**<br>(HB3**(157)**:FCB1**(192)**) | **FC27**<br>(K1**(407)**:HB3**(335)**) | **3D7**<br>(3D7**(264)**:FCB1**(341)**) |
|---|---|---|---|---|
| Run 1 | ≥3000:1 | 1:20 | 1000:1 | 1:100 |
| Run 2 | 100:1 | 1:20 | 100:1 | 1:50 |

**Supplementary Table 8.** Detection limit and robustness of both runs of high-resolution capillary electrophoresis using *msp1* and *msp2* with **small fragment minority** applying the 10% cut-off. For each marker the amplicon size of each strain is indicated in bp.

**LIMIT OF DETECTION FOR DIFFERENT ALLELIC FAMILIES**

| | **K1**<br>(3D7**(247)**:K1**(177)**) | **Mad20**<br>(HB3**(157)**:FCB1**(192)**) | **FC27**<br>(K1**(407)**:HB3**(335)**) | **3D7**<br>(3D7**(264)**:FCB1**(341)**) |
|---|---|---|---|---|
| Runs identical | ≥3000:1 | ≥1:3000 | ≥3000:1 | 1:500 |

**Supplementary Table 9.** Detection limit and robustness of both runs of high-resolution capillary electrophoresis using microsatellite markers with **short fragment in minority** applying the microsatellite cut-offs of Greenhouse et al., 2006. For each marker the amplicon size of each strain is indicated in bp (3D7:K1:HB3:FCB1).

**LIMIT OF DETECTION FOR MICROSATELLITES**

| | ***Pf*PK2**<br>(**247**:**247**:270:241) | **TA40**<br>(270:**249**:255:**249**) | **TA60**<br>(325:**322**:319:**322**) | **TA81**<br>(182:188:191:179) |
|---|---|---|---|---|

| | | | | |
|---|---|---|---|---|
| Runs identical | 20:20:20:1 | ≥1500:1:1500:1 | ≥1000:1000:1:1000 | ≥1500:1500:1500:1 |

***msp1 and msp2 genotyping using high-resolution melting analysis.*** The *msp1* products of *P. falciparum* strains 3D7 and FCB1 as well as K1 and HB3 shared almost identical melting temperatures, which resulted in an overlap in mixed-strains sample peaks in the ratio 1:1:1:1 (Supplementary Figure 2). This impeded the discrimination between the two strains in all of the samples.

*Msp2* PCR products of the strains 3D7 and FCB1 produced two melt peaks. The *msp2* product of the strains 3D7, K1 and FCB1 shared almost identical $T_m$, except for the second $T_m$ for 3D7 and FCB1 *msp2* product. However, the second $T_m$ peak was lower than the first peak of each strain. For all the low parasitaemia samples (10 p/µl), no $T_m$ was detected and therefore the samples resulted in a negative outcome. In all samples (10 p/µl and 1000 p/µl), a $T_m$ was observed at around 80.85 °C (Figure 2).

Because the *msp2* products of 3D7 (peak 1), K1 and FCB1 (peak 1) shared almost identical $T_m$ an overlap of mixed-strains sample peaks in a ratio of 1:1:1:1 was observed (Figure). Peak 2 of 3D7 and FCB1 *msp2* product was not detected in this sample. This made it difficult to distinguish between each of the three strains. In addition, it was questionable if the presence of HB3 could really be excluded in this sample with the threshold or if the peak was overlapping for all the four strains and only shifted right because the other three strains' *msp2* product possessed a higher $T_m$ (Figure 2).

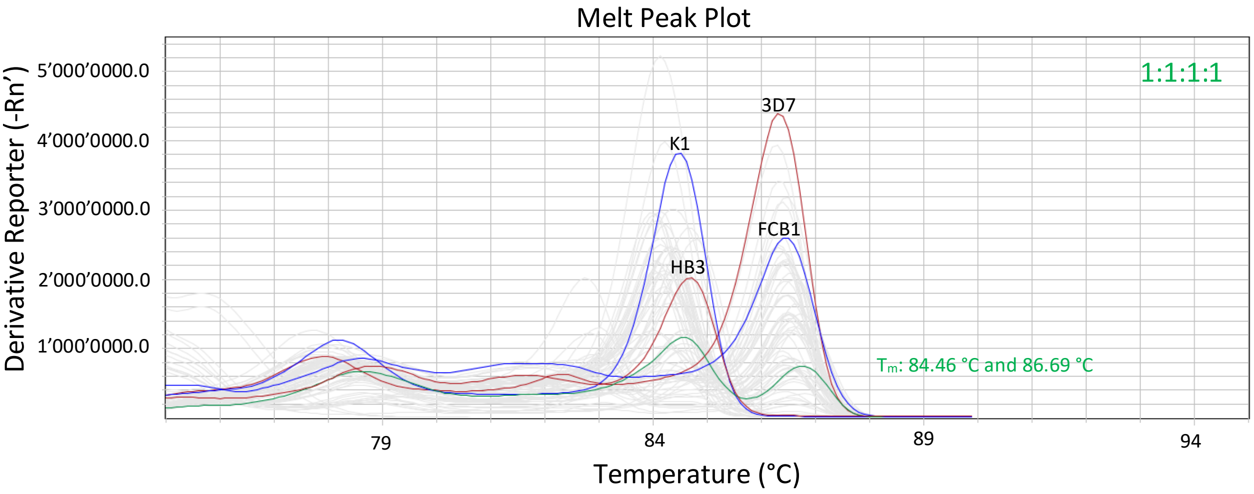

**Supplementary Figure 2.** Melt peak plot from one of the four single-strain sample peaks (*msp1*) (1000 p/µl) in red and blue as well as the 1:1:1:1 [3D7:K1:HB3:FCB1] mixed-strains sample (10 p/µl each strain) with mean melting temperatures ($T_m$) in green.

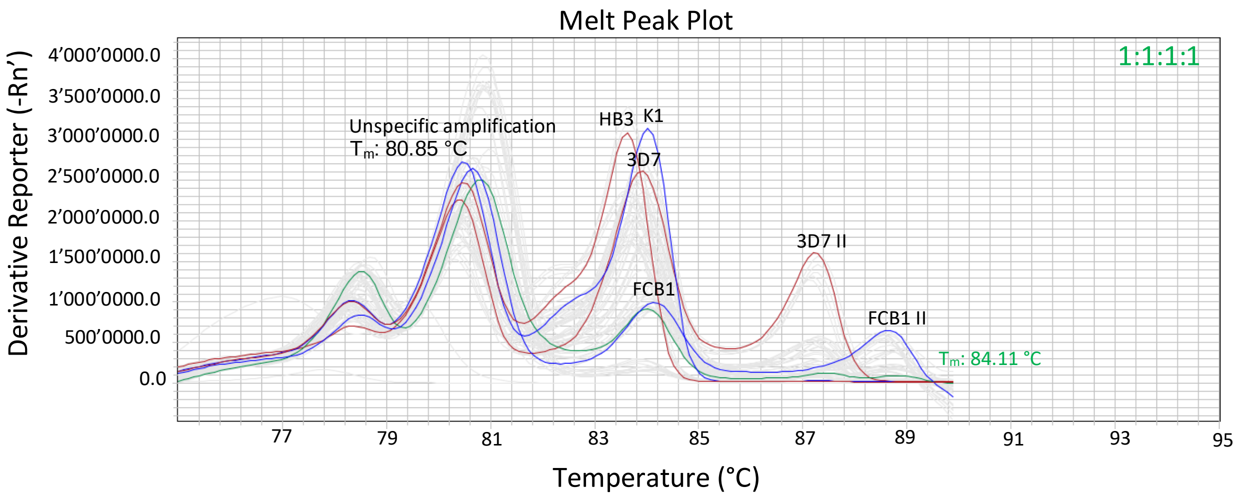

**Supplementary Figure 3.** Melt peak plot from one of the four single-strain sample peaks (*msp2*) (1000 p/µl) in red and blue as well as the 1:1:1:1 [3D7:K1:HB3:FCB1] mixed-strains sample (10 p/µl each strain) with mean melting temperature ($T_m$) in green.

**Supplementary Table 10.** a) HRM *msp1* results form run 1 and run 2 using the 0.2 °C threshold. Box shading color: orange = this $T_m$ belongs to 3D7 strain; yellow = K1; green = HB3; turquoise = FCB1; pink-purple = 3D7 and/or FCB1; light blue = K1 and/or HB3; gray box shading color in the ratio colon indicates a second plate due to lack of space in a 96-well plate for all the samples. Black font color = temperature clearly belongs to one strain or two strains when threshold is applied; red font color = $T_m$ does not belong to any strain if threshold is applied. The orange font color in combination with a specific box shading color of a certain strain shows that with the threshold applied it would belong to the box shading color strain but another strain cannot be excluded on the basis of the ratio. Pos. rep. = positive replicate; av. = average; $T_m$ = melting temperature. The SD deviation of each triplicate of a sample is shown. b) Another HRM was performed with different single-strain sample concentrations. The SD deviation of each triplicate of a sample is shown.

a)

| | Run 1 | | | | Run 2 | | | |
|---|---|---|---|---|---|---|---|---|
| Ratio<br><br>3D7:K1:HB3:FCB1 | 3D7 and/or FCB1 of pos. rep. | 3D7 and/or FCB1 av. $T_m$ (°C) of triplicate | K1 and/or HB3 of pos. rep. | K1 and/or HB3 av. $T_m$ (°C) of triplicate | 3D7 and/or FCB1 of pos. rep. | 3D7 and/or FCB1 av. $T_m$ (°C) of triplicate | K1 and/or HB3 of pos. rep. | K1 and/or HB3 av. $T_m$ (°C) of triplicate |
| 1:0:0:0 (10 p/µl) | 100 | 86.56 ± 0.09 | | | 100 | 86.59 ± 0.06 | | |
| 1:0:0:0 (1000 p/µl) | 100 | 86.32 ± 0.06 | | | 100 | 86.44 ± 0.08 | | |
| 0:1:0:0 (10 p/µl) | | | 100 | 84.49 ± 0.03 | | | 100 | 84.49 ± 0.06 |
| 0:1:0:0 (1000 p/µl) | | | 100 | 84.27 ± 0.18 | | | 100 | 84.32 ± 0.05 |
| 0:0:1:0 (10 p/µl) | | | 100 | 84.90 ± 0.06 | | | 100 | 84.85 ± 0.08 |
| 0:0:1:0 (1000 p/µl) | | | 100 | 84.70 ± 0.06 | | | 100 | 84.77 ± 0.09 |
| 0:0:0:1 (10 p/µl) | 100 | 86.56 ± 0.05 | | | 100 | 86.54 ± 0.03 | | |
| 0:0:0:1 (1000 p/µl) | 100 | 86.49 ± 0.06 | | | 100 | 86.61 ± 0.05 | | |
| 1:1:1:1 | 100 | 86.69 ± 0.11 | 100 | 84.46 ± 0.20 | 100 | 86.72 ± 0.10 | 100 | 84.56 ± 0.08 |
| 5:1:1:1 | 100 | 86.52 ± 0.10 | 100 | 84.27 ± 0.23 | 100 | 86.59 ± 0.08 | 100 | 84.29 ± 0.19 |
| 1:5:1:5 | 100 | 86.59 ± 0.10 | 100 | 84.24 ± 0.10 | 100 | 86.62 ± 0.08 | 100 | 84.39 ± 0.08 |
| 1:5:5:5 | 100 | 86.66 ± 0.13 | 100 | 84.37 ± 0.22 | 100 | 86.67 ± 0.08 | 100 | 84.46 ± 0.06 |
| 1:10:1:1 | 0 | | 100 | 84.46 ± 0.14 | 0 | | 100 | 84.47 ± 0.05 |
| 1:1:10:10 | 100 | 86.67 ± 0.10 | 100 | 84.49 ± 0.19 | 100 | 86.67 ± 0.10 | 100 | 84.51 ± 0.11 |
| 10:1:10:10 | 100 | 86.41 ± 0.09 | 100 | 84.12 ± 0.13 | 100 | 86.46 ± 0.05 | 100 | 84.29 ± 0.03 |
| 1:1:15:1 | 0 | | 100 | 84.54 ± 0.15 | 0 | | 100 | 84.74 ± 0.03 |
| 15:15:1:1 | 100 | 86.69 ± 0.08 | 100 | 84.41 ± 0.12 | 100 | 86.60 ± 0.05 | 100 | 84.42 ± 0.09 |
| 15:15:1:15 | 100 | 86.44 ± 0.03 | 100 | 84.06 ± 0.03 | 100 | 86.51 ± 0.00 | 100 | 84.31 ± 0.06 |
| 1:1:1:20 | 100 | 86.54 ± 0.06 | 33 | 84.27 | 100 | 86.54 ± 0.06 | 100 | 84.29 ± 0.03 |
| 20:1:20:1 | 100 | 86.42 ± 0.10 | 100 | 84.46 ± 0.25 | 100 | 86.44 ± 0.03 | 100 | 84.54 ± 0.06 |
| 20:20:20:1 | 100 | 86.39 ± 0.03 | 100 | 84.16 ± 0.06 | 100 | 86.37 ± 0.08 | 100 | 84.24 ± 0.08 |
| 50:1:1:1 | 100 | 86.39 ± 0.08 | 0 | | 100 | 86.42 ± 0.03 | 0 | |
| 1:50:1:50 | 100 | 86.52 ± 0.08 | 100 | 84.09 ± 0.16 | 100 | 86.46 ± 0.05 | 100 | 84.13 ± 0.05 |
| 1:50:50:50 | 100 | 86.47 ± 0.03 | 100 | 84.14 ± 0.08 | 100 | 86.47 ± 0.03 | 100 | 84.16 ± 0.03 |
| 1:0:0:0 (10 p/µl) | 100 | 86.24 ± 0.08 | | | 100 | 86.75 ± 0.05 | | |
| 1:0:0:0 (1000 p/µl) | 100 | 86.22 ± 0.03 | | | 100 | 86.67 ± 0.06 | | |
| 0:1:0:0 (10 p/µl) | | | 100 | 84.18 ± 0.00 | | | 100 | 84.69 ± 0.03 |

| | | | | | | | | |
|---|---|---|---|---|---|---|---|---|
| 0:1:0:0 (1000 p/µl) | | | 100 | 84.21 ± 0.06 | | | 100 | 84.77 ± 0.10 |
| 0:0:1:0 (10 p/µl) | | | 100 | 84.67 ± 0.03 | | | 100 | 85.15 ± 0.03 |
| 0:0:1:0 (1000 p/µl) | | | 100 | 84.60 ± 0.06 | | | 100 | 85.12 ± 0.09 |
| 0:0:0:1 (10 p/µl) | 100 | 86.31 ± 0.00 | | | 100 | 86.75 ± 0.09 | | |
| 0:0:0:1 (1000 p/µl) | 100 | 86.34 ± 0.03 | | | 100 | 86.75 ± 0.05 | | |
| 100:1:1:1 | 100 | 86.36 ± 0.05 | 0 | | 100 | 86.84 ± 0.10 | 0 | |
| 1:100:1:100 | 100 | 86.42 ± 0.03 | 100 | 84.22 ± 0.05 | 100 | 86.89 ± 0.03 | 100 | 84.60 ± 0.06 |
| 1:1000:1000:1000 | 100 | 86.48 ± 0.04 | 100 | 84.34 ± 0.08 | 100 | 87.05 ± 0.13 | 100 | 84.77 ± 0.13 |
| 1:500:1:1 | 0 | | 100 | 84.34 ± 0.03 | 0 | | 100 | 84.75 ± 0.06 |
| 1:1:1500:500 | 100 | 86.37 ± 0.03 | 100 | 84.39 ± 0.03 | 100 | 86.87 ± 0.03 | 100 | 84.75 ± 0.03 |
| 500:1:500:500 | 100 | 86.36 ± 0.09 | 100 | 84.40 ± 0.11 | 100 | 86.82 ± 0.03 | 100 | 84.79 ± 0.03 |
| 1:1:1000:1 | 0 | | 100 | 84.65 ± 0.03 | 0 | | 100 | 85.10 ± 0.06 |
| 1000:1000:1:1 | 100 | 86.24 ± 0.03 | 100 | 84.16 ± 0.03 | 100 | 86.85 ± 0.00 | 100 | 84.69 ± 0.03 |
| 1000:1000:1:1000 | 100 | 86.37 ± 0.03 | 100 | 84.19 ± 0.03 | 100 | 86.82 ± 0.08 | 100 | 84.59 ± 0.08 |
| 1:1:1:1500 | 100 | 86.28 ± 0.03 | 0 | | 100 | 86.66 ± 0.00 | 0 | |
| 1500:1:1500:1 | 100 | 86.24 ± 0.06 | 100 | 84.56 ± 0.06 | 100 | 86.65 ± 0.05 | 100 | 84.90 ± 0.03 |
| 1500:1500:1500:1 | 100 | 86.23 ± 0.06 | 100 | 84.24 ± 0.06 | 100 | 86.62 ± 0.06 | 100 | 84.59 ± 0.06 |

b)

| Ratio | of pos. rep. | $T_m$ |
|---|---|---|
| 1:0:0:0 (100 p/µl) | 100 | 86.36 ± 0.00 |
| 1:0:0:0 (500 p/µl) | 100 | 86.36 ± 0.07 |
| 0:1:0:0 (100 p/µl) | 100 | 84.36 ± 0.06 |
| 0:1:0:0 (500 p/µl) | 100 | 84.47 ± 0.05 |
| 0:0:1:0 (100 p/µl) | 100 | 84.82 ± 0.05 |
| 0:0:1:0 (500 p/µl) | 100 | 84.80 ± 0.08 |
| 0:0:0:1 (100 p/µl) | 100 | 86.42 ± 0.08 |
| 0:0:0:1 (500 p/µl) | 100 | 86.54 ± 0.03 |

**Supplementary Table 11.** a) HRM *msp2* results from run 1 and run 2 using the 0.2 °C threshold. Box shading color: orange = this $T_m$ belongs to 3D7 strain; yellow = K1; green = HB3; turquoise = FCB1; pink-purple = 3D7 and/or FCB1; light blue = K1 and/or HB3; gray box shading color in the ratio colon indicates a second plate due to lack of space in a 96-well plate for all the samples. Black font color = temperature clearly belongs to one strain or two strains when threshold is applied (depending on the box shading color); red font color = $T_m$ does not belong to any strain if threshold is applied; purple font color = temperature can belong to either 2/3 or 2/4 or 3/4 strains. The orange font color in combination with a specific box shading color of a certain strain, shows that with the threshold applied it would belong to the box shading color strain but another strain cannot be excluded on the basis of the ratio. Pos. rep. = positive replicate; av. = average; $T_m$ = melting temperature. The SD deviation of each triplicate of a sample is shown. b) Another HRM was performed with different single-strain sample concentrations. The SD deviation of each triplicate of a sample is shown.

a)

| | Run 1 | | | | Run 2 | | | |
|---|---|---|---|---|---|---|---|---|
| Ratio 3D7:K1:HB3:FCB1 | 3D7 and/or K1 and/or FCB1 of pos. rep. | 3D7 and/or K1 and/or FCB1 av. $T_m$ (°C) of triplicate | All strains possible of pos. rep. | All strains possible av. $T_m$ (°C) of triplicate | 3D7 and/or K1 and/or FCB1 of pos. rep. | 3D7 and/or K1 and/or FCB1 av. $T_m$ (°C) of triplicate | All strains possible of pos. rep. | All strains possible av. $T_m$ (°C) of triplicate |
| 1:0:0:0 (10 p/µl) | 0 | | 0 | | 0 | | 0 | |
| 1:0:0:0 (1000 p/µl) | 100 | 83.91 ± 0.03 | 100 | 83.91 ± 0.03 | 100 | 83.84 ± 0.2 | 100 | 83.84 ± 0.2 |
| | | 87.23 ± 0.03 | | 87.23 ± 0.03 | | 87.17 ± 0.16 | | 87.17 ± 0.16 |
| 0:1:0:0 (10 p/µl) | 0 | | 0 | | 0 | | 0 | |
| 0:1:0:0 (1000 p/µl) | 100 | 84.02 ± 0.02 | 100 | 84.02 ± 0.02 | 100 | 84.19 ± 0.11 | 100 | 84.19 ± 0.11 |
| 0:0:1:0 (10 p/µl) | | | 0 | | | | 0 | |
| 0:0:1:0 (1000 p/µl) | | | 100 | 83.64 ± 0.03 | | | 100 | 83.51 ± 0.06 |
| 0:0:0:1 (10 p/µl) | 0 | | 0 | | 0 | | 0 | |
| 0:0:0:1 (1000 p/µl) | 100 | 84.07 ± 0.09 | 100 | 84.07 ± 0.09 | 100 | 84.09 ± 0.03 | 100 | 84.09 ± 0.03 |
| | | 88.56 ± 0.08 | | 88.56 ± 0.08 | | 88.57 ± 0.03 | | 88.57 ± 0.03 |
| 1:1:1:1 | 100 | 84.11 ± 0.08 | | | 100 | 84.06 ± 0.10 | | |
| 5:1:1:1 | 100 | 84.01 ± 0.03 | | | 100 | 83.96 ± 0.08 | | |
| | | 87.35 ± 0.00 | | | | 87.27 ± 0.04 | | |
| 1:5:1:5 | 100 | 84.16 ± 0.14 | | | 100 | 84.11 ± 0.08 | | |
| | | | | | | 88.74 ± 0.14 | | |

| | | | | | | | | |
|---|---|---|---|---|---|---|---|---|
| 1:5:5:5 | 100 | 84.99 ± 0.03 | | | 100 | 83.94 ± 0.08<br>88.69 ± 0.00 | | |
| 1:10:1:1 | 100 | 84.07 ± 0.00 | | | 100 | 84.04 ± 0.06 | | |
| 1:1:10:10 | | | 100 | 83.79 ± 0.12<br>88.85 ± 0.12 | | | 100 | 83.63 ± 0.09<br>88.74 ± 0.10 |
| 10:1:10:10 | | | 100 | 83.68 ± 0.05<br>87.28 ± 0.06 | | | 100 | 83.63 ± 0.05<br>87.27 ± 0.03 |
| 1:1:15:1 | | | 100 | 83.69 ± 0.03 | | | 100 | 83.66 ± 0.03 |
| 15:15:1:1 | 100 | 84.07 ± 0.05<br>87.36 ± 0.03 | | | 100 | 83.98 ± 0.05<br>87.32 ± 0.03 | | |
| 15:15:1:15 | 100 | 83.98 ± 0.05 | | | 100 | 83.96 ± 0.08<br>87.20 ± 0.05 | | |
| 1:1:1:20 | 100 | 84.01 ± 0.10<br>88.67 ± 0.06 | | | 100 | 84.01 ± 0.08<br>88.69 ± 0.10 | | |
| 20:1:20:1 | | | 100 | 83.63 ± 0.05<br>87.26 ± 0.08 | | | 100 | 83.48 ± 0.00<br>87.15 ± 0.05 |
| 20:20:20:1 | | | 100 | 83.76 ± 0.03<br>87.15 ± 0.00 | | | 100 | 83.66 ± 0.03<br>87.07 ± 0.03 |
| 50:1:1:1 | 100 | 83.86 ± 0.08<br>87.25 ± 0.09 | | | 100 | 83.75 ± 0.06<br>87.10 ± 0.00 | | |
| 1:50:1:50 | | | 100 | 83.84 ± 0.03<br>88.57 ± 0.06 | 100 | 83.75 ± 0.03<br>88.49 ± 0.05 | | |
| 1:50:50:50 | | | 100 | 83.61 ± 0.03<br>88.54 ± 0.00 | | | 100 | 83.51 ± 0.03<br>88.47 ± 0.03 |
| 1:0:0:0 (10 p/µl) | 0 | | 0 | | 0 | | 0 | |
| 1:0:0:0 (1000 p/µl) | 100 | 83.63 ± 0.00<br>86.92 ± 0.03 | 100 | 83.63 ± 0.00<br>86.92 ± 0.03 | 100 | 83.58 ± 0.05<br>86.85 ± 0.09 | 100 | 83.58 ± 0.05<br>86.85 ± 0.09 |
| 0:1:0:0 (10 p/µl) | 0 | | 0 | | 0 | | 0 | |
| 0:1:0:0 (1000 p/µl) | 100 | 83.84 ± 0.03 | 100 | 83.84 ± 0.03 | 100 | 83.78 ± 0.05 | 100 | 83.78 ± 0.05 |
| 0:0:1:0 (10 p/µl) | | | 0 | | | | 0 | |
| 0:0:1:0 (1000 p/µl) | | | 100 | 83.38 ± 0.09 | | | 100 | 83.36 ± 0.06 |

| | | | | | | | | |
|---|---|---|---|---|---|---|---|---|
| 0:0:0:1 (10 p/µl) | 0 | | 0 | | 0 | | 0 | |
| 0:0:0:1 (1000 p/µl) | 100 | 83.96 ± 0.03<br>88.46 ± 0.03 | 100 | 83.96 ± 0.03<br>88.46 ± 0.03 | 100 | 83.91 ± 0.03<br>88.40 ± 0.03 | 100 | 83.91 ± 0.03<br>88.40 ± 0.03 |
| 100:1:1:1 | 100 | 83.73 ± 0.00<br>87.22 ± 0.03 | | | 100 | 83.64 ± 0.03<br>87.15 ± 0.05 | | |
| 1:100:1:100 | 100 | 83.71 ± 0.03<br>88.54 ± 0.08 | | | 100 | 83.64 ± 0.03<br>88.49 ± 0.00 | | |
| 1:1000:1000:1000 | | | 100 | 83.50 ± 0.08<br>88.57 ± 0.10 | | | 100 | 83.43± 0.05<br>88.55 ± 0.08 |
| 1:500:1:1 | 100 | 83.83 ± 0.05 | | | 100 | 83.78 ± 0.05 | | |
| 1:1:1500:500 | | | 100 | 83.13 ± 0.00<br>88.46 ± 0.03 | | | 100 | 83.12 ± 0.03<br>88.44 ± 0.00 |
| 500:1:500:500 | | | 100 | 83.20 ± 0.00<br>87.05 ± 0.10 | | | 100 | 83.17 ± 0.08<br>87.03 ± 0.13 |
| 1:1:1000:1 | | | 100 | 83.33 ± 0.05 | | | 100 | 83.32 ± 0.06 |
| 1000:1000:1:1 | | | 100 | 83.49 ± 0.03<br>86.99 ± 0.06 | | | 100 | 83.43 ± 0.05<br>86.97 ± 0.08 |
| 1000:1000:1:1000 | | | 100 | 83.56 ± 0.03<br>87.08 ± 0.03 | | | 100 | 83.50 ± 0.03<br>87.00 ± 0.05 |
| 1:1:1:1500 | 100 | 83.73 ± 0.05<br>88.36 ± 0.03 | | | 100 | 83.71 ± 0.10<br>88.36 ± 0.11 | | |
| 1500:1:1500:1 | | | 100 | 83.07 ± 0.03<br>86.95 ± 0.05 | | | 100 | 83.08 ± 0.09<br>86.97 ± 0.10 |
| 1500:1500:1500:1 | | | 100 | 83.37 ± 0.03<br>86.97 ± 0.06 | | | 100 | 83.25 ± 0.03<br>86.93 ± 0.06 |

b)

| Ratio | of pos. rep. | $T_m$ |
|---|---|---|
| 1:0:0:0 (100 p/µl) | 100 | 83.81 ± 0.03<br>86.90 ± 0.05 |

| | | |
|---|---|---|
| 1:0:0:0 (500 p/µl) | 100 | 83.58 ± 0.05 |
| | | 86.70 ± 0.05 |
| 0:1:0:0 (100 p/µl) | 100 | 83.78 ± 0.05 |
| 0:1:0:0 (500 p/µl) | 100 | 83.55 ± 0.08 |
| 0:0:1:0 (100 p/µl) | 100 | 83.35 ± 0.08 |
| 0:0:1:0 (500 p/µl) | 100 | 83.13 ± 0.09 |
| 0:0:0:1 (100 p/µl) | 100 | 83.86 ± 0.03 |
| | | 88.24 ± 0.05 |
| 0:0:0:1 (500 p/µl) | 100 | 83.67 ± 0.05 |
| | | 88.09 ± 0.05 |

Applying the WHO algorithm for NI/R determination, in only 2/20 samples all methods agree on NI and in 10/20 samples all methods agree on R. Thus, in 8/20 samples there is discrepancy between the different genotyping methods (Table 9).

**Table 12.** Comparison of new infection/recrudescence outcome of different genotyping methods using the WHO algorithm.

| **Sample ID** | **F-CE**: *msp1, msp2, glurp* | **HRM**: *msp1, msp2* | **H-CE**: *msp1, msp2, glurp* | **H-CE**: 4 microsatellites | **H-CE**: *msp1, msp2* and 4 microsatellites | **TADS**: *ama1, cpmp, csp, cpp, msp7* |
|---|---|---|---|---|---|---|
| **S1** | R | R | R | R | R | R |
| **S10** | R | R | R | R | R | R |
| **S11** | R | R | R | R | R | R |
| **S12** | R | R | R | R | R | R |
| **S13** | R | R | R | R | R | R |
| **S14** | NI | R | R | R | R | R |
| **S15** | NI | R | R | NI | NI | NI |
| **S16** | R | R | R | R | R | R |
| **S17** | R | R | NI | R | NI | R |
| **S18** | NI | NI | NI | R | NI | NI |
| **S19** | R | NI | R | R | R | R |
| **S2** | NI | NI | NI | R | R | R |
| **S20** | NI | NI | NI | NI | NI | NI |
| **S3** | R | R | R | R | R | R |
| **S4** | R | NI | R | R | R | R |
| **S5** | R | R | R | R | R | R |

| | | | | | | |
|---|---|---|---|---|---|---|
| **S6** | NI | NI | NI | NI | NI | NI |
| **S7** | NI | R | R | R | R | R |
| **S8** | R | R | R | R | R | R |
| **S9** | R | R | R | R | R | R |

Applying the 2/3 algorithm for NI/R determination, in none of the samples all methods agree on NI and in 17/20 samples all methods agree on R. Thus, in 3/20 samples there is discrepancy between the different genotyping methods. However, using the 2/3 algorithm the outcome tends to be R compared to the WHO algorithm (Table 10).

**Table 13.** Comparison of new infection/recrudescence outcome of different genotyping methods using the 2/3 algorithm (HRM not available since only two markers were used, for H-CE microsatellites a 3/4 and 4/6 algorithm was used when combined with *msp1* and *msp2*, for TADS a 3/5 algorithm was used).

| **Sample ID** | **F-CE**: *msp1, msp2, glurp* | **H-CE**: *msp1, msp2, glurp* | **H-CE**: microsatellites | **H-CE**: *msp1, msp2* and 4 microsatellites | **TADS**: ama1, cpmp, csp, cpp, msp7 |
|---|---|---|---|---|---|
| **S1** | R | R | R | R | R |
| **S10** | R | R | R | R | R |
| **S11** | R | R | R | R | R |
| **S12** | R | R | R | R | R |
| **S13** | R | R | R | R | R |
| **S14** | R | R | R | R | R |
| **S15** | R | R | R | R | NI |
| **S16** | R | R | R | R | R |
| **S17** | R | R | R | R | R |
| **S18** | R | NI | R | R | NI |
| **S19** | R | R | R | R | R |
| **S2** | R | R | R | R | R |
| **S20** | R | NI | R | NI | NI |

| | | | | | |
|---|---|---|---|---|---|
| **S3** | R | R | R | R | R |
| **S4** | R | R | R | R | R |
| **S5** | R | R | R | R | R |
| **S6** | NI | R | NI | NI | NI |
| **S7** | R | R | R | R | R |
| **S8** | R | R | R | R | R |
| **S9** | R | R | R | R | R |

***For F-CE (msp1, msp2 and glurp)***, in 13/20 samples all markers agree on R and in none of the samples all the markers show a NI outcome simultaneously. Thus, 7/20 samples show discordant results. The outcome of the two algorithms differ, using the WHO algorithm 7/20 NI outcomes are counted whereas for the 2/3 algorithm only 1/20 NI outcome is counted.

**Supplementary Table 14.** Comparison of new infection/recrudescence outcome of different markers using fast capillary electrophoresis including the difference in outcomes when using two algorithms.

| **Outcome** | **Count** | **F-CE marker results** | | | **Classification** | |
|---|---|---|---|---|---|---|
| | | *msp1* | *msp2* | *glurp* | WHO algorithm | 2/3 algorithm |
| Clear recrudescence | 13 | R | R | R | R | R |
| Clear new infection | 0 | NI | NI | NI | NI | NI |
| Intermediate results | 3 | R | R | NI | NI | R |
| | 3 | R | NI | R | NI | R |
| | 1 | NI | NI | R | NI | NI |
| **Final outcome** | | | | | | |
| Recrudescence | | | | | 13 | 19 |
| New infection | | | | | 7 | 1 |
| Total | | | | | 20 | 20 |

***For H-CE (msp1, msp2 and glurp)***, in 15/20 samples all markers agree on R and in none of the samples all the markers show a NI outcome simultaneously. Thus, 5/20 samples show discordant results. The outcome of the two algorithms differ, using the WHO algorithm 5/20 NI outcomes are counted whereas for the 2/3 algorithm only 2/20 NI outcome is counted.

**Supplementary Table 15.** Comparison of new infection/recrudescence outcome of different markers using high-resolution capillary electrophoresis including the difference in outcomes when using two algorithms.

| **Outcome** | **Count** | **H-CE marker results** | | | **Classification** | |
|---|---|---|---|---|---|---|
| | | *msp1* | *msp2* | *glurp* | WHO algorithm | 2/3 algorithm |
| Clear recrudescence | 15 | R | R | R | R | R |
| Clear new infection | 0 | NI | NI | NI | NI | NI |
| Intermediate results | 2 | R | R | NI | NI | R |
| | 1 | R | NI | R | NI | R |
| | 1 | NI | R | NI | NI | NI |
| | 1 | R | NI | NI | NI | NI |
| **Final outcome** | | | | | | |
| Recrudescence | | | | | 15 | 18 |
| New infection | | | | | 5 | 2 |
| Total | | | | | 20 | 20 |

***For H-CE (microsatellites)***, in 17/20 samples all markers agree on R and in none of the samples all the markers show a NI outcome simultaneously. Thus, 3/20 samples show discordant results. The outcome of the two algorithms differ, using the WHO algorithm 3/20 NI outcomes are counted whereas for the 2/3 algorithm 2/20 NI outcome is counted.

**Supplementary Table 16.** Comparison of new infection/recrudescence outcome of different markers using high-resolution capillary electrophoresis including the difference in outcomes when using two algorithms.

| Outcome | Count | H-CE marker results | | | | Classification | |
|---|---|---|---|---|---|---|---|
| | | *Pf*PK2 | TA40 | TA60 | TA81 | WHO algorithm | 3/4 algorithm |
| Clear recrudescence | 18 | R | R | R | R | R | R |
| Clear new infection | 0 | NI | NI | NI | NI | NI | NI |
| Intermediate results | 1 | R | R | R | NI | NI | R |
| | 2 | NI | NI | NI | R | NI | NI |
| **Final outcome** | | | | | | | |
| Recrudescence | | | | | | 17 | 18 |
| New infection | | | | | | 3 | 2 |
| Total | | | | | | 20 | 20 |

***Comparison of results when microsatellites replace glurp. For H-CE (msp1, msp2 and either PfPK2, TA40 or TA60)***, in 16/20 samples all markers agree on R and in none of the samples all the markers show a NI outcome simultaneously. Thus, 4/20 samples show discordant results. The outcome of the two algorithms differ, using the WHO algorithm 4/20 NI outcomes are counted whereas for the 2/3 algorithm only 1/20 NI outcome is counted.

**Supplementary Table 17 a.** Comparison of new infection/recrudescence outcome of different markers using high-resolution capillary electrophoresis including the difference in outcomes when using two algorithms. The combination of *msp1/msp2* and either microsatellites *Pf*PK2, TA40 or TA60 is shown.

| Outcome | Count | H-CE marker results | | | Classification | |
|---|---|---|---|---|---|---|
| | | *msp1* | *msp2* | *Pf*PK2, TA40 or TA60 | WHO algorithm | 2/3 algorithm |
| Clear recrudescence | 16 | R | R | R | R | R |
| Clear new infection | 0 | NI | NI | NI | NI | NI |
| Intermediate results | 1 | R | R | NI | NI | R |
| | 2 | R | NI | R | NI | R |

| | | | | | | |
|---|---|---|---|---|---|---|
| | 1 | NI | R | NI | NI | NI |
| **Final outcome** | | | | | | |
| Recrudescence | | | | | 16 | 19 |
| New infection | | | | | 4 | 1 |
| Total | | | | | 20 | 20 |

***For H-CE (msp1, msp2 andTA81)***, in 16/20 samples all markers agree on R and in none of the samples all the markers show a NI outcome simultaneously. Thus, 4/20 samples show discordant results. The outcome of the two algorithms differ, using the WHO algorithm 3/20 NI outcomes are counted whereas for the 2/3 algorithm no NI outcome is counted.

**Supplementary Table 17 b.** Comparison of new infection/recrudescence outcome of different markers using high-resolution capillary electrophoresis including the difference in outcomes when using two algorithms. The combination of *msp1/msp2* and microsatellite TA81 is shown.

| **Outcome** | **Count** | **H-CE marker results** | | | **Classification** | |
|---|---|---|---|---|---|---|
| | | *msp1* | *msp2* | TA81 | WHO algorithm | 2/3 algorithm |
| Clear recrudescence | 16 | R | R | R | R | R |
| Clear new infection | 0 | NI | NI | NI | NI | NI |
| Intermediate results | 1 | NI | R | R | NI | R |
| | 2 | R | NI | R | NI | R |
| | 1 | R | R | NI | NI | R |
| **Final outcome** | | | | | | |
| Recrudescence | | | | | 16 | 20 |
| New infection | | | | | 4 | 0 |
| Total | | | | | 20 | 20 |

***For HRM (msp1 and msp2)***, in 14/20 samples all markers agree on R and in 2/20 samples all the markers show a NI outcome simultaneously. Thus, 4/20 samples show discordant results. Only the WHO algorithm could be applied since only two markers were used for this method.

**Supplementary Table 18.** Comparison of new infection/recrudescence outcome of different markers using high-resolution melting.

| **Outcome** | **Count** | **HRM marker results** | | **Classification** |
|---|---|---|---|---|
| | | *msp1* | *msp2* | WHO algorithm |
| Clear recrudescence | 14 | R | R | R |
| Clear new infection | 2 | NI | NI | NI |
| Intermediate results | 2 | R | NI | NI |
| | 2 | NI | R | NI |
| **Final outcome** | | | | |
| Recrudescence | | | | 14 |
| New infection | | | | 6 |
| Total | | | | 20 |

***For amplicon deep sequencing (ama1-D3, cpmp, csp, cpp and msp7),*** in 16/20 samples all markers agree on R and in four samples all the markers agree on a NI outcome. Thus, 1/20 samples show a discordant result. The outcome of the two algorithms do not differ.

**Supplementary Table 19.** Comparison of new infection/recrudescence outcome of different markers using amplicon deep sequencing including the difference in outcomes when using two algorithms.

| **Outcome** | **Count** | **TADS marker results** | | | | | **Classification** | |
|---|---|---|---|---|---|---|---|---|
| | | *ama1* | *cpmp* | *csp* | *cpp* | *msp7* | WHO algorithm | 3/5 algorithm |
| Clear recrudescence | 16 | R | R | R | R | R | R | R |
| Clear new infection | 3 | NI | NI | NI | NI | NI | NI | NI |

| | | | | | | | | |
|---|---|---|---|---|---|---|---|---|
| Intermediate results | 1 | NI | NI | NI | NI | R | NI | NI |
| **Final outcome** | | | | | | | | |
| Recrudescence | | | | | | | 16 | 16 |
| New infection | | | | | | | 4 | 4 |
| Total | | | | | | | 20 | 20 |

#### Supplementary Methods

##### 1. Fast Capillary electrophoresis (QIAxcel)

Protocol adapted from for PCR [15] and [27] for the analysis.

Workflow of amplicon fragment length analysis by fast-CE.

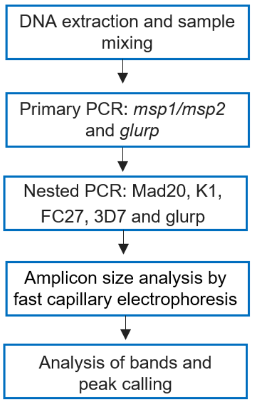

All the PCRs were performed using the thermocycler Biometra Tone 96 Gradient.

<u>Primary PCR of *msp1*, *msp2* and *glurp*</u>

Master-mix preparations for multiplex primary PCR targeting *msp1* and *msp2* of *P. falciparum*.

| Reagent | Working solution | Final concentration | per 1 reaction (µl) |
|---|---|---|---|
| $ddH_2O$ | | | 28 |
| 10x Buffer B | 10x | 1x | 5 |
| dNTPs | 2 mM | 200 µM | 5 |
| $MgCl_2$ | 25 mM | 2 mM | 4 |
| Primer Mix: M1-OF, M1-OR, M2-OF, M2-OR | 10 µM | 0.5 µM | 2.5 |
| **FIRE Pol®** Polymerase | 5 U/µl | 0.05 U/µl | 0.5 |
| DNA solution | | | 5 |
| **Total volume** | | | **50** |

All the reagents, including the DNA solution, were vortexed and quickly spun down, except for the FIRE Pol® Polymerase, which was spun down and mixed by pipetting up and down before added into the mix. Before the plate was transferred into the thermocycler it was spun down for approx. 10 seconds at 1200 RPM.

Thermocycling conditions for multiplex primary PCR targeting *msp1* and *msp2* of *P. falciparum*.

| Temperature | Time | Cycles |
|---|---|---|
| 94°C | 05:00 min | 1x |
| 94°C | 00:30 min | 30x |
| 54°C | 01:00 min | 30x |
| 72ºC | 01:00 min | 30x |
| 72°C | 05:00min | 1x |
| 4°C | ∞ | 1x |

The master-mix preparations for singleplex primary PCR targeting *glurp* of *P. falciparum*, is the same as for multiplex primary PCR targeting *msp1* and *msp2* except for the primer mix: G4 & G5mod (10 µM).

Thermocycling conditions for singleplex primary PCR targeting *glurp* of *P. falciparum.*

| Temperature | Time | Cycles |
|---|---|---|
| 94°C | 05:00 min | 1x |
| 94°C | 01:00 min | 30x |
| 50°C | 01:00 min | 30x |

| | | |
|---|---|---|
| 72°C | 02:00 min | 30x |
| 72°C | 10:00min | 1x |
| 4°C | ∞ | 1x |

<u>Nested PCR of *msp1* allelic families: Mad20 and K1</u>

Master-mix preparations for singleplex nested PCR targeting *msp1* Mad20 allelic family of *P. falciparum*.

| Reagent | Working solution | Final concentration | per 1 reaction (µl) |
|---|---|---|---|
| $ddH_2O$ | | | 32.5 |
| 10x Buffer B | 10x | 1x | 5 |
| dNTPs | 2 mM | 200 µM | 5 |
| $MgCl_2$ | 25 mM | 2 mM | 4 |
| Primer Mix: M1-MF/M1-MR | 10 µM | 0.4 µM | 2 |
| **FIRE Pol®** Polymerase | 5 U/µl | 0.05 U/µl | 0.5 |
| *msp1/msp2* pPCR product | | | 1 |
| **Total volume** | | | **50** |

Thermocycling conditions for singleplex nested PCR targeting *msp1* Mad20 allelic family of *P. falciparum*.

| Temperature | Time | Cycles |
|---|---|---|
| 94°C | 05:00 min | 1x |

| | | |
|---|---|---|
| 94°C | 00:30 min | 30x |
| 59°C | 01:00 min | 30x |
| 72°C | 01:00 min | 30x |
| 72°C | 05:00min | 1x |
| 4°C | ∞ | 1x |

For singleplex nested PCR targeting K1, the master-mix contained the equal amount of identical reagents as for Mad20 singleplex nested PCR except for the primer mix where the primer pair M1-KF/M1-KR (10 µM) was used. The thermocycler conditions for K1 singleplex nested PCR were identical to Mad20 singleplex nested PCR.

<u>Nested PCR of *msp2* allelic families: FC27 and 3D7</u>

Master-mix preparations for singleplex nested PCR targeting *msp2* FC27 allelic family of *P. falciparum.*

| Reagent | Working solution | Final concentration | per 1 reaction (µl) |
|---|---|---|---|
| $ddH_2O$ | | | 33.5 |
| 10x Buffer B | 10x | 1x | 5 |
| dNTPs | 2 mM | 200 µM | 5 |
| $MgCl_2$ | 25 mM | 1.5 mM | 3 |
| Primer Mix: Stail - M5 | 10 µM | 0.4 µM | 2 |
| **FIRE Pol®** Polymerase | 5 U/µl | 0.05 U/µl | 0.5 |
| *msp1/msp2* pPCR product | | | 1 |
| **Total volume** | | | **50** |

Thermocycling conditions for singleplex nested PCR targeting *msp2* FC27 allelic family of *P. falciparum*.

| Temperature | Time | Cycles |
|---|---|---|
| 94°C | 05:00 min | 1x |
| 94°C | 00:30 min | 30x |
| 50°C | 00:45 min | 30x |
| 72°C | 01:00 min | 30x |
| 72°C | 10:00min | 1x |
| 4°C | ∞ | 1x |

For singleplex nested PCR targeting 3D7, the master-mix contained the equal amount of identical reagents as for FC27 singleplex nested PCR except for the primer pair Stail - N5 (10 µM) was added. The thermocycler conditions were identical to FC27 singleplex nested PCR.

<u>Nested PCR of *glurp*</u>

The master-mix preparations for singleplex nested PCR targeting *glurp* of *P. falciparum* is the same as for singleplex nested PCR targeting 3D7 and FC27 except for the primer Mix: GNF-G3 and *glurp* pPCR product was used as a template

Thermocycling conditions for singleplex nested PCR targeting *glurp* of *P. falciparum*.

| Temperature | Time | Cycles |
|---|---|---|
| 94°C | 05:00 min | 1x |
| 94°C | 01:00 min | 30x |
| 58°C | 01:00 min | 30x |

| | | |
|---|---|---|
| 72°C | 01:00 min | 30x |
| 72°C | 10:00min | 1x |
| 4°C | ∞ | 1x |

<u>Fast capillary electrophoresis</u>

The High-resolution cartridge was used and run with the OM500 program for medium DNA concentration. Everything was prepared according to the protocol provided in the QIAxcel manual user [27]. Briefly 10 µl of nested PCR product was transferred into a new 96 PCR well plate and empty wells were filled with 10 µl QX DNA dilution buffer to have full rows of liquid. The PCR plates where the samples were taken from needed to be covered by aluminum foil to protect the light sensitive samples. After the sample transfer, the plates were put back to the fridge for later use in high-resolution capillary electrophoresis, the second method. After adding all the samples and the buffer the plate was spun down for 1 minute at 4000 RPM remove the bubbles on the bottom. Samples were run with an alignment and size marker. For the alignment marker (15 bp/3 kb), 12 wells of a multiply®-µStrip 0.2 ml chain was filled with 15 µl alignment marker and a drop of mineral oil in each well and was renewed every week. For the size marker (100 bp-2.5 kb), 10 µl of 1:10 diluted size marker was added and needed to be refilled only if necessary. The run was around 12 minutes for 12 samples (one row). After the run, the gel images and electropherograms were analyzed.

<u>Data analysis of fast-CE</u>

The samples were analyzed using the integrated QIAxcel ScreenGel software of the fast capillary electrophoresis machine. The expected fragment size variation is 10 bp for a 100-500 bp fragments (expected *msp* size) and for a 500-1000 bp fragments (expected *glurp* size) the variation may reach up to 50 bp [27]. Thus, the threshold for each sample to have a present strain or not depended on the fragment size to be within the ± 10 bp for *msp* and ± 50 bp for *glurp*. To apply this, the rounded mean of the positive control triplicates in each plate was calculated. Then the variation e.g. for *msp,* 10 bp was divided by the mean of the positive control and the outcome was multiplied by 100 to get the tolerance in percent. For instance, K1 plate 1, the mean of positive control triplicate of 3D7 strain equaled 254 bp. Next, 10 (the variation for *msp*) was divided by the mean 254 and multiplied by 100, which equaled 3.9%. Thus, all the fragment sizes in this plate, which equaled 254 bp ± 3.9% were tolerated as a peak of interest. This threshold for each strain in each plate was calculated because the fragment size estimation in the positive controls slightly differed between the plates. The integrated QIAxcel ScreenGel software enabled "peak calling" where all the peaks within this specific tolerance are found and listed in a "peak calling table". This table allowed fast and easy analysis of each ratio showing the peak of interest or n/a (not applicable) where no peak within the tolerance was found, meaning the strain was not detected. This information from the peak calling table was then summarized in an Excel table showing how many replicates in percent where positive for which ratio. The columns of the table were divided in allelic families as well as if the longer or shorter fragment within this family is in minority. This allowed the direct comparison of the detection limit between the different markers as well as the difference it makes if the short or long fragment is in minority. The table also enabled the comparison of the different runs revealing the robustness of the method. For NI/R analysis, strains from D0 and DX of each sample were considered the same when within the threshold range described above (± 10 bp for *msp* and ± 50 bp for *glurp*) and considered different strains when out of threshold range.

#### 2. High-Resolution Capillary electrophoresis: *msp1, msp2* and *glurp*

Protocol adapted from [15].

Workflow of high-resolution CE.

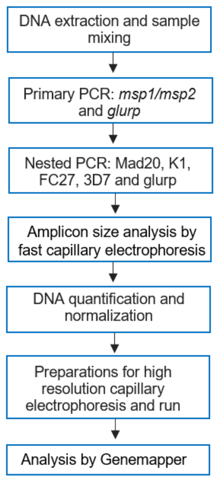

The same markers were used as described in methods above. Since the primary and nested PCR were already done for method 1 (F-CE), the first step for the second method started with the DNA concentration measurement of the nested PCR products.

<u>DNA quantification</u>

For estimating the DNA concentration (ng/µl) of the nested PCR products, the SYNERGY H1 microplate reader from BioTek® and the Promega QuantiFluor® dsDNA System kit - and protocol was used [28]. Briefly, for the master-mix the 20x TE buffer needed to be diluted 1:20 with MiliQ water. Next, the dsDNA dye needed to be diluted 1:400 with 1x TE buffer. 200 µl of the 1x TE buffer/dsDNA dye mix was added per well in a black corning 96 well multi-well plate and mixed with either 1 µl nested PCR product or 10 µl of standard. A standard was used in duplicate: 20 ng/µl, 5 ng/µl, 1.25 ng/µl, 0.31 ng/µl, 0.078 ng/µl, 0.02 ng/µl, 0.005 ng/µl and 1x TE buffer (0 ng/µl). With the software Gen5 3.08, the fluorescence intensity information was collected and exported into an Excel file, where this output was converted into DNA concentration (ng/µl).

Sample normalization using a liquid handling robot

Target concentration needed for high-resolution capillary electrophoresis.

| **Marker** | **Target concentration** |
|---|---|
| Mad20 | 3 – 4 ng/µl |
| K1 | 1.5 – 2.5 ng/µl |
| FC27 | 0.6 – 1.5 ng/µl |
| 3D7 | 0.8 – 1.5 ng/µl |
| *Glurp* | 2 – 3 ng/µl |

Due to high sample numbers, the sample normalization the samples was performed using a liquid handling robot (QIAgility from Qiagen). The estimated DNA concertation (ng/µl) from the nested PCR products measured by the microplate reader were transferred to the software of the liquid handling robot (software version 4.17.1). Then the normalization concentrations from the above table, were set as target concentration depending on the marker. With this given information, automated pipetting for sample normalization could be performed with MiliQ water.

High-resolution capillary electrophoresis preparation

Next a mix containing 12 µl Hi-Di™ Formamide and 0.2 µl GeneScan™ - 500 LIZ® size standard (for all *msp* allelic families) or 1200 LIZ® (*glurp*) size standard was prepared for one reaction. 12 µl of this mix was used per sample and 2 µl of the normalized nested PCR product was then added to the PCR well. Empty wells were filled with 14 µl MiliQ water, so all wells of the MicroAmp® Optical 96-Well reaction plate with barcode were filled with liquid.

The plates were subsequently sent to Microsynth for the high-resolution capillary electrophoresis run. There the samples were first denatured for 3 minutes at 95 °C and cooled down to 4 °C before performing high-resolution capillary electrophoresis. The samples were run on the ABI3730XL capillary electrophoresis machine from Thermo Fisher.

<u>Data analysis of high-resolution capillary electrophoresis</u>

The peaks from high-resolution capillary electrophoresis were analysed by GeneMapper® Software 6 from Applied Biosystems.

All samples were analyzed by checking the presence of the fragment of interest and its RFU (peak height in electropherogram). Two cut-offs were used to assess the detection limit:

i) <u>The ≥500 RFU:</u> If in 2 out of 3 replicates the minority clones showed <500 RFU the sample peaks were considered as background. (Usually used to assess if a peak is background for instance in negative template controls (NTC)).

ii) <u>The 10/20% cut-off:</u> If in 2 out of 3 replicates the minority clone's fragment is smaller than 10% of the highest peak (*msp1/msp2*) or smaller than 20% of the highest peak (*glurp*) the sample was considered as negative. (The 10/20% cut-off is applied in genotyping working with patient's blood samples to ensure real peaks are picked for the analysis.)

This information was summarized in an Excel table, showing the percentage of positive replicates for each ratio. First, the columns were divided in each marker and the allelic families were further divided in short or long fragment minority and those colons were again divided in ≥500 RFU and 10/20% threshold. Thus, this table allowed the direct comparison of:

i) the two thresholds within one plate (difference in detection limit between the thresholds)

ii) of short and long fragment minority (difference in detection limit between fragment length)

iii) the different markers used (difference in detection limit between the markers)

iv) the two runs (difference in detection limit of repeated experiment revealing the robustness of the method)

For NI/R outcome, peaks in electropherograms of D0 and DX of each sample were compared. Strains were considered the same if peaks (with 500 RFU and 10/20% cut-off applied) have the same size with variation up to 2bp.

##### 3. High-Resolution Capillary electrophoresis: microsatellites

Protocol adapted from [9], small changes were made to optimally work in our laboratory (see words written in bold-type).

<u>Single-round PCR of *Pf*PK2, TA40, TA60 and TA81 microsatellites</u>

Master-mix preparations for single-round and simplex PCR targeting *Pf*PK2, TA40, TA60 or TA81 microsatellites markers of *P. falciparum.*

| Reagent | Working solution | Final concentration | per 1 reaction (µl) |
|---|---|---|---|
| $ddH_2O$ | | | **17.3** |
| Buffer II | 10x | 1x | **2.5** |
| dNTPs each | 2 mM | 40 µM | **0.5** |
| $MgCl_2$ | 25 mM | 1.5 mM | **1.5** |
| Primer each fw/rv | 10 µM | 200 nmol/l | **0.5** |
| **AmpliTap Gold DNA Polymerase** | 5 U/µl | 0.04 U/µl | **0.2** |
| Template | | | **2.5** |
| **Total volume** | | | **25** |

Thermocycling conditions for single-round PCR targeting for *Pf*PK2, TA40, TA60 and TA81 of *P. falciparum*.

| **Thermocycling conditions: PfPK2** | | **Thermocycling conditions: TA40** | | **Thermocycling conditions: TA60** | | **Thermocycling conditions: TA81** | |
|---|---|---|---|---|---|---|---|
| 95ºC - 10:00 min | | 95ºC - 10:00 min | | 95ºC - 10:00 min | | 95ºC - 10:00 min | |
| 95ºC - 00:15 min | | 95ºC - 00:15 min | | 95ºC - 00:15 min | | 95ºC - 00:15 min | |
| 52.3ºC - 01:30 min | 40 cycles | 53 ºC - 01:30 min | 40 cycles | 53 ºC - 01:30 min | 40 cycles | 58.9 ºC - 01:30 min | 40 cycles |
| 60ºC - 00:30 min | | 60ºC - 00:45 min | | 60ºC - 00:30 min | | 60ºC - 00:30 min | |
| 60ºC - 05:00 min | | 60ºC - 05:00 min | | 60ºC - 05:00 min | | 60ºC - 05:00 min | |
| Then go to 4°C ∞ | | Then go to 4°C ∞ | | Then go to 4°C ∞ | | Then go to 4°C ∞ | |

Fast capillary electrophoresis

Fast capillary electrophoresis was performed as described in method 1 to check if amplification of the different marker worked.

DNA quantification and Sample normalization using a liquid handling robot

DNA quantification and sample normalization was performed as described in method 2. The final concentration of each microsatellite was:

<table>
<tr><th>Marker</th><th>Target concentration</th></tr>
<tr><td>PfPK2</td><td rowspan="3">0.8 ng/µl</td></tr>
<tr><td>TA40</td></tr>
<tr><td>TA60</td></tr>
<tr><td>TA81</td><td>1.2 ng/µl</td></tr>
</table>

High-resolution capillary electrophoresis preparation

Next a mix containing 10 µl Hi-Di™ Formamide and 0.2 µl GeneScan™ - 500 LIZ® size standard (for all microsatellites) was prepared for one reaction. 10 µl of this mix was used per sample and 2 µl of the normalized PCR product was then added to the PCR well. Empty wells were filled with 12 µl MiliQ water, so all wells of the MicroAmp® Optical 96-Well reaction plate with barcode were filled with liquid.

The plates were subsequently sent to Microsynth for the high-resolution capillary electrophoresis run. There the samples were first denatured for 3 minutes at 95 °C and cooled down to 4 °C before performing high-resolution capillary electrophoresis. The samples were run on the ABI3730XL capillary electrophoresis machine from Thermo Fisher.

Data analysis of high-resolution capillary electrophoresis

The peaks from high-resolution capillary electrophoresis were analysed by GeneMapper® Software 6 from Applied Biosystems using the cut-off from [9].

The script used for microsatellite cut-off analysis can be found under the following link: https://github.com/SwissTPH/AmpSeqAnalysis.

**4. Amplicon Deep Sequencing**

Protocol adapted from [13,22,29], small changes were made to optimally work in our laboratory (see words written in bold-type).

Workflow of targeted Amp-Seq.

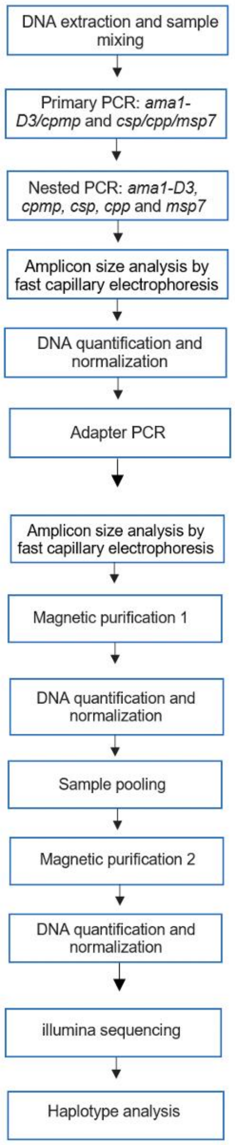

The markers used for targeted Amp-Seq

Overview of the markers used for targeted Amp-Seq and their fragment size (3D7 strain), which is the same for all strains. For the nested PCR amplicons, the forward and reverse linker sequences are included (22 bp and 23 bp). For the adapter PCR products forward and reverse adapter sequences (67 bp and 68 bp) are included.

| | **Amplicon size in bp** | | |
|---|---|---|---|
| **Marker** | **Primary PCR** | **Nested PCR** | **Adapter PCR** |
| *ama1-D3* | 588 | 559 | 689 |
| *cpmp* | 601 | 473 | 603 |
| *csp* | 478 | 372 | 502 |
| *cpp* | 392 | 438 | 568 |
| *msp7* | 457 | 419 | 549 |

Primary duplex and triplex PCR of *ama1-D3/cpmp* and *csp/cpp/msp7*

Note: All reaction preparations containing KAPA HiFi HotStart Ready mix should be prepared on ice.

Master-mix preparations for duplex primary PCR targeting *ama1-D3* and *cpmp* of *P. falciparum.*

| Reagent | Working solution | Final concentration | per 1 reaction (µl) |
|---|---|---|---|
| $H_2O$ | | | 3.75 |
| KAPA HiFi HotStart Ready mix | 2x | 1x | 7.5 |
| Primer pair 1 (fw/rv) | 10 µM | 250 nm | 0.375 |
| Primer pair 2 (fw/rv) | 10 µM | 250 nm | 0.375 |

| | | | |
|---|---|---|---|
| Primer pair 3 (fw/rv) | 10 µM | 250 nm | 0 |
| DNA template | | | 3 |
| **Total volume** | | | **15** |

All the reagents for the master-mix were first vortexed and quickly spun down, before added into the mix. Next, DNA solution was vortexed and 3 µl added to the master-mix, making a total of 15 µl per reaction. Before the plate was transferred into the thermocycler it was spun down for approx. 10 seconds at 1200 RPM.

Master-mix preparations for triplex primary PCR targeting *csp, cpp* and *msp7* of *P. falciparum.*

| Reagent | Working solution | Final concentration | per 1 reaction (µl) |
|---|---|---|---|
| $H_2O$ | | | 3.375 |
| KAPA HiFi HotStart Ready mix | 2x | 1x | 7.5 |
| Primer pair 1 (fw/rv) | 10 µM | 250 nm | 0.375 |
| Primer pair 2 (fw/rv) | 10 µM | 250 nm | 0.375 |
| Primer pair 3 (fw/rv) | 10 µM | 250 nm | 0.375 |
| DNA template | | | 3 |
| **Total volume** | | | **15** |

Thermocycling conditions for duplex primary PCR targeting *ama1-D3* and *cpmp* and triplex primary PCR targeting *csp, cpp* and *msp7* of *P. falciparum.*

| Temperature | Time | Cycles |
|---|---|---|
| 95°C | 03:00 min | 1x |
| 98°C | 00:20 min | 20x |
| 54°C* | 00:15 min | 20x |
| 72°C | 00:45 min | 20x |
| 72°C | 02:00min | 1x |
| 4°C | ∞ | 1x |

*52 °C for master-mix preparations for triplex primary PCR targeting *csp, cpp* and *msp7*

Nested singleplex PCR of *ama1-D3, cpmp, csp, cpp* or *msp7*

Master-mix preparations for simplex nested PCR targeting *ama1-D3, cpmp, csp, cpp* or *msp7* of *P. falciparum*.

| Reagent | Working solution | Final concentration | per 1 reaction (µl) |
|---|---|---|---|
| $H_2O$ | | | 5.5 |
| KAPA HiFi HotStart Ready mix | 2x | 1x | 10 |
| Primer pair (fw/rv) | 10 µM | 250 nm | 0.5 |
| DNA template | | | 4 |
| **Total volume** | | | **20** |

Thermocycling conditions for simplex nested PCR targeting *ama1-D3, cpmp, csp, cpp* or *msp7* of *P. falciparum*.

| Temperature | Time | Cycles |
|---|---|---|

| | | |
|---|---|---|
| 95°C | 03:00 min | 1x |
| 98°C | 00:20 min | 10x |
| 55°C | 00:15 min | 10x |
| 72°C | 00:45 min | 10x |
| 98°C | 00:20min | 10x |
| 62°C | 00:15 min | 10x |
| 72°C | 00:45 min | 10x |
| 72°C | 01:30 min | 1x |
| 4°C | ∞ | 1x |

The master-mix content and thermocycler conditions for *cpmp* needed to be optimized due to low PCR product concentration compared to the other markers.

Master-mix preparations for efficiency optimization simplex nested PCR targeting *cpmp* of *P. falciparum.*

| Reagent | Working solution | Final concentration | per 1 reaction (µl) |
|---|---|---|---|
| $H_2O$ | | | **4.5** |
| KAPA HiFi HotStart Ready mix | 2x | 1x | 10 |
| Primer pair (fw/rv) | 10 µM | 250 nm | 0.5 |
| DNA template | | | **5** |

| | | | |
|---|---|---|---|
| **Total volume** | | | **20** |

Thermocycling conditions for efficiency optimization simplex nested PCR targeting *cpmp* of *P. falciparum.*

| Temperature | Time | Cycles |
|---|---|---|
| 95°C | 03:00 min | 1x |
| 98°C | 00:20 min | 10x |
| 55°C | 00:15 min | 10x |
| 72°C | 00:45 min | 10x |
| 98°C | 00:20min | **15x** |
| 62°C | 00:15 min | **15x** |
| 72°C | 00:45 min | **15x** |
| 72°C | 01:30 min | 1x |
| 4°C | ∞ | 1x |

Fast capillary electrophoresis: DNA Fast Analysis Cartridge

The QIAxcel DNA Fast Analysis Cartridge was used to check the bands after the nested PCR, using the protocol from Qiagen [27]. A total of 10 µl was loaded into QIAxcel, 4 µl of nPCR product and 6 µl of water to have enough nPCR product for the adapter PCR.

DNA quantification

The same protocol for the microplate reader was used as described in 2.9.1, except for 2 µl DNA was added instead of 1 µl. The following standard was used in duplicate: 20 ng/µl, 10 ng/µl, 5 ng/µl, 1.25 ng/µl, 0.31 ng/µl, 0.078 ng/µl, 0.02 ng/ µl and 1x TE buffer (0 ng/µl).

Sample normalization using a liquid handling robot

Due to high sample numbers, the pipetting for normalizing the samples was performed using a liquid handling robot (used in this project QIAgility from Qiagen). All the samples were normalized to a concentration of 10 ng/µl if possible. For the samples having a starting concentration of 5 ng/µl to <10 ng/µl were normalized to the end concentration of 5 ng/µl. Samples with <1 ng/µl starting concentration as well as the negative controls were used undiluted. The samples were diluted using the Ambion™ Nucleasse-Free water.

Adapter duplex and triplex PCR of *ama1-D3/cpmp* and *csp/cpp/msp7*

The normalized plates were first spun down quickly. A new PCR plate was prepared to mix 4 µl of each marker *csp, cpp* and *msp7*. In another PCR plate, 6 µl of *cpmp* and 4 µl of *ama1-D3* was mixed. Those mixed samples were the template DNA for the next PCR.

Master-mix preparations for adapter PCR.

| Reagent | Working solution | Final concentration | per 1 reaction (µl) |
|---|---|---|---|
| $H_2O$ | | | 2.5 |
| KAPA HiFi HotStart Ready Mix | 2x | 1x | 7.5 |
| Primer fw | 5 µM | 333 nm | 1 |
| Primer rev | 5 µM | 333 nm | 1 |
| DNA template mix | | | **3** |
| **Total volume** | | | **15** |

First, water and the KAPA Ready Mix were mixed and added in each well. The forward and reverse primers were first added in a separate multiply®-µStrip 0.2 ml chain to ease the transfer. Following 1 µl each forward and reverse primer was added separately in each well containing the rest of the master-mix. In the end the DNA was added.

Thermocycling conditions for adapter PCR.

| Temperature | Time | Cycles |
|---|---|---|
| 95°C | 03:00 min | 1x |
| 98°C | 00:20 min | 10x |
| 58°C | 00:30 min | 10x |
| 72°C | 00:45 min | 10x |
| 72°C | 02:00 min | 1x |
| 4°C | ∞ | 1x |

Magnetic purification

To clean up and size-select the samples, magnetic purification was performed. The protocol was used from NucleoMag® NGS Clean-up and Size Select, which was sent with the magnetic beads. The magnetic beads were removed from the fridge to bring to room temperature and were then vortexed until it was a homogenous solution. Next, 8 µl of bead solution and 10 µl of adapter PCR template was added in each well of a 96-well 300 µl round bottom plate to achieve a ratio of 0.8. The beads were mixed with the template by pipetting 10 times up and down. To separate the magnetic beads from the solution, the plate was placed on a 96-well magnetic separation system and after 5 minutes at room temperature, the solution appeared clear and the magnetic beads were attracted to the sides on the bottom. The supernatant was removed without disturbing the magnetic beads. It was important to work fast to avoid that the magnetic beads dry out. The plate stayed on the magnetic separation system while washing each well with 200 µl of 80% ethanol. Next, the plate was incubated at room temperature for 30 seconds and the supernatant was removed. To dry the beads, the plate, still on the magnetic separation system, was incubated at room temperature for 5-15 minutes for complete evaporation of the remaining alcohol. For the elution of the target DNA , the 96 -well plate was removed from the magnetic stand and 15 µl of elution buffer (1x TE buffer) was added in each well and pipetted up and down 10 times to re-suspend the magnetic beads. The plate was incubated for 2-5 minutes and then was placed on the magnetic stand again for at least 5 minutes until the liquid was clear. Then 15 µl of purified DNA was transferred in a new 96- well PCR plate.

DNA quantification

The concentration of the purified adapter PCR products was measured using the same machine, protocol, standard and kit as described above.

Second normalization (manually)

The samples were normalized by manual pipetting because due to the duplex and triplex adapter PCR there were less samples compared to the number of samples after the singleplex nested PCR. Thus, the manual normalization would have taken the same amount of time. The measured purified adapter PCR products were diluted manually to the concentration of around 25 ng/µl, which corresponds to around 45 nM. The final concentration to send for sequencing is 20 nM and due to another magnetic purification and thus possible loss of DNA, this normalization was set to 45 nM. The samples were diluted using the Ambion™ Nucleasse-Free water.

Sample pooling

The adapter PCR primers bearing a specific 8 nucleotide molecular barcode allows the pooling (multiplexing) and later de-multiplexing of the samples [29]. For pooling the different samples and negative controls together, 4 µl of each sample was transferred to a 1.5 ml Eppendorf tube. The plates of *ama1-D3/cpmp* run 1 and run 2 were pooled separately in one tube. The same was done for *csp/cpp/msp7*.

Second magnetic purification

For the final magnetic purification the same protocol as described in 2.10.8 was used except that 100 µl DNA template of each of the pooled samples was used. Therefore, to get a ratio of 0.8, 80 µl of beads was used. Due to the high volume, the washing step was done twice and the DNA was eluted in 40 µl elution buffer (1x TE buffer).

Qubit Fluorometer

To measure the DNA concentration of the eight pooled tubes, Qubit was performed using the Qubit 3 Fluorometer (dsDNA Broad range) and Qubit™ dsDNA BR Assay Kit. For this, two times 190 µl Qubit® working solution was prepared consisting of 1:200 diluted Qubit® dsDNA BR Reagent in Qubit® dsDNA BR Buffer. Subsequently 10 µl of each of the standards (standard 1 with 0 ng/µl and standard 2 with 5 ng/µl) was added in two Qubit™ assay tubes, vortexed, spun down and measured before the samples, to set the standard. For the samples, 198 µl of Qubit® working solution for each sample and 2 µl of sample DNA was mixed in Qubit™ assay tubes, vortexed, spun down and then measured [30].

Fast capillary electrophoresis: DNA High-resolution Cartridge

The High-resolution Cartridge was used due to high concentration and long fragment size of the amplicons to get more accurate band size estimations. It was performed in the same way as described above.

Final manual normalization

To dilute the eight samples to the desired concentration of 20 nM, the Qubit results were used for the calculations (measured DNA conc. in ng/µl). The formula was found in [31]. The 660 g/mol corresponds to the molecular weight of DNA.

Formula 1: The mean of the base pair length of *ama1-D3* and *cpmp* (646 bp) was used:

$$\frac{Measured\ DNA\ conc.in\ ng/\mu l}{660\ g/mol\ \times 646\ bp} \times 10^{6} = conc.in\ nM$$

Formula 2: The mean of the base pair length of *cpp, csp* and *msp7* (539.6) was used:

$$\frac{Measured\ DNA\ conc.in\ ng/\mu l}{660\ g/mol\ \times 539.6\ bp} \times 10^{6} = conc.in\ nM$$

Formula 3: The output from the previous formula was used as c1; c2 = 20 nM, the desired end concentration; v2 = 40 µl, the desired end volume. The output of this formula shows the amount of DNA in µl that will be used for the normalization and this number minus 40 µl equals the amount of TE buffer added:

$$c1 \times v1 = c2 \times v2$$

$$v1 = \frac{c2 \times v2}{c1}$$

Next, 40 µl of the pooled *ama1-D3/cpmp* were transferred in a new tube. The same was performed for pooled *cpp/csp/msp7*. The markers *ama1-D3/cpmp* and *cpp/csp/msp7* were still separated. In the end, two Eppendorf tubes containing 40 µl of pooled samples were sent for sequencing by the MiSeq illumina instrument at the University hospital Basel. Paired-end mode using the MiSeq reagent kit v3 (600 cycles) was used and run with 15% spike-in of Enterobacteria phage *phiX* control v3 [13].

<u>Data analysis of targeted Amp-Seq</u>

About 10'000 reads coverage was targeted. The stored base call data was converted into sequence data as FASTQ files [32]. FASTQ files were retrieved from the Scicore server at the University of Basel. The data were accessed via MobaXterm software [22]. FastQC was run for quality control of the data and then the data was analysed in R using the R package HaplotypR [22]. The clusters were de-multiplexed by sample possessing an identical index and by marker possessing the same nested PCR primer sequence [22,32]. Then the primer sequence was truncated to increase sequence quality and the forward and reverse reads were merged [22,29]. The merged reads were then aligned to 3D7 reference strain and called for SNPs: >50% mismatch rate to call the SNP a genotype and ≥2 samples have this SNP [22,29]. Then the reads were called for haplotypes (a sequence variant of an amplicon [29]): At least 3 reads should have identical sequences in ≥2 samples, ≥0.1% within-sample haplotype frequency (a certain haplotype in a sample needs to be present in a frequency of at least 0.1% of total haplotypes present in the sample) is required [29]. If only one read showed a specific haplotype (singleton) in a sample, it was considered as background [22]. Chimeric reads (introduced by incomplete primer extension followed by in-homologous re-annealing), indels (introduced by polymerase slippage) or noise (samples with a sequence coverage less than 25 reads) were considered as PCR artefacts [22]. The number of reads per haplotype per sample were summarized in Excel from which the proportion of total reads, including PCR artefacts, contaminant haplotypes (real haplotypes not detected in positive controls) as well as background, was calculated.

The scripts used for amplicon deep sequencing haplotype calling, can be found under the following link: https://github.com/SwissTPH/AmpSeqAnalysis.

#### 5. High-resolution melt analysis (HRM)

Protocol adapted from [12], small changes were made to optimally work in our laboratory (see words written in bold-type).

Workflow of HRM.

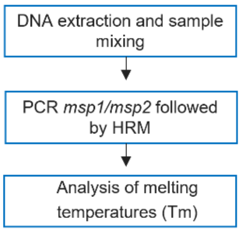

<u>The markers used for high-resolution melting</u>

Overview of the markers used and their corresponding fragment size and GC-content [33].

| **Marker** | ***msp1*** | | ***msp2*** | |
|---|---|---|---|---|
| **Strain** | **Fragment size (bp)** | **GC-content (%)** | **Fragment size (bp)** | **GC-content (%)** |
| 3D7 | 321 | 42.4 | 549 | 42.1 |
| K1 | 285 | 40.7 | not available | not available |
| HB3 | 240 | 40 | 501 | 38.5 |
| FCB1 | 276 | 42 | not available | not available |

<u>Final and adapted PCR-HRM targeting *msp1* and *msp2*</u>

Adapted and final master-mix preparations for quantitative PCR targeting *msp1* and *msp2* of *P. falciparum* based on [12]. The bold text indicates where changes were made compared to the original protocol from [12].

| Reagent | Working solution | Final concentration | per 1 reaction (µl) |
|---|---|---|---|

| | | | |
|---|---|---|---|
| $H_2O$ | | | 7.55 |
| $NH_4$ | 10x | 1x | 2.5 |
| dNTPs | 2mM each | **300 µM each*** | 3.75 |
| $MgCl_2$ | 50 mM | 3 mM | 1.5 |
| Primer forward | 10 µM | **400 nM*** | 1 |
| Primer reverse | 10 µM | **400 nM*** | 1 |
| Taq Polymerase | 5 U/µl | 1 U | 0.2 |
| Syto Green | 50 µM | 5 µM | 2.5 |
| DNA template | | | 5 |
| **Total volume** | | | **25** |

*For protocol optimization, the dNTP concentration was increased from 300 nM to 300 µM and the primer concentration was increased from 200 nM to 400 nM (final concentration in master-mix).

All the reagents including the DNA solution were vortexed and spun down before added to the master-mix except for the Tag polymerase, which was spun down first followed by softly pipetting up and down. The plate needed to be centrifuged before putting into the machine. The PCR plate was covered with MicroAmp™ Optical Adhesive Film, rather than 8-strip opt. clear flat caps for accurate fluorescence measurement by the machine. The machine used for the PCR and later HRM step was QuantStudio5 from Applied Biosystems.

Adapted and final thermocycling conditions for quantitative PCR targeting *msp1* and *msp2* of *P. falciparum* [12].

| Temperature | Time | Cycles |
|---|---|---|

| | | |
|---|---|---|
| 94°C | 02:00 min | 1x |
| 94°C | 00:15 min | **45x*** |
| 54°C | 00:20 min | **45x*** |
| 72°C | 00:40 min | **45x*** |

After each cycle, the fluorescence intensity was measured. This was followed by the HRM with the following conditions: ramping 0.1 °C/**5 seconds*** between 75 °C and 90 °C.

* For protocol optimization, the PCR cycle number was altered from 40 to 45, and the melting step was changed from 0.1 °C/1 second to 0.1 °C/5 seconds between 75-90 °C.

Data analysis of high-resolution melting

For analysing the melting temperatures ($T_m$) of each sample, QuantStudio™ Design and Analysis v1.5.1 software from Applied Biosystems as well as Excel was used. With the QuantStudio™ Design and Analysis software a melt peak plot of each sample and any sample combination could be visualized. This enabled the direct comparison of $T_m$ peaks of different ratios as well as the visualization of overlapping peaks. The collected data was transferred on Excel. There the mean $T_m$ of each triplicate was then calculated. The standard deviation (SD) of the positive controls on each plate was calculated as well. In this method, also a threshold was applied: if a $T_m$ differs by ≤0.2 °C from a $T_m$ in another sample, it is considered as the same strain [12]. Thus, if a $T_m$ differs by >0.2 °C from a $T_m$ in another sample, it is considered as two distinct strains. Therefore, knowing the $T_m$ from the single-strain samples ($T_m$ from each strain) enabled the peak assignment to the present strains of each multi-strain sample. Each plate has new positive controls, from which $T_m$ were used to assign the $T_m$ from multi-strain samples in this plate. For the NI/R outcome $T_m$ was compared from D0 and DX of each sample.

**Primer Sequences for all markers used in this study**

Method 1 and 2: Fast capillary electrophoresis and high-resolution capillary electrophoresis

Primer sequences of *P. falciparum* multiplex primary PCR targeting *msp1* and *msp2*

| Name | Sequence |
|---|---|
| M1-OF (*msp1* outer forward) | 5’-CTAGAAGCTTTAGAAGATGCAGTATTG-3’ |
| M1-OR | 5’-CTTAAATAGTATTCTAATTCAAGTGGATCA-3’ |

| Name | Sequence |
|---|---|
| M2-OF (*msp2* outer forward) | 5’-ATGAAGGTAATTAAAACATTGTTCTCTATTATA-3’ |
| M2-OR | 5’-CTTTGTTACCATCGGTACATTCTT-3’ |

Primer sequences for *P. falciparum* primary PCR targeting *glurp*

| Name | Sequence |
|---|---|
| G4_fw | 5’-ACATGCAAGTGTTGATCC-3’ |
| G5mod_rev | 5’-CAGATGGTTTGGGAGTAACGTT-3’ |

Primer sequences for *P. falciparum* nested PCR targeting *msp1* Mad20

| Name | Sequence |
|---|---|
| M1-MF | Tail 5’-AAATGAAGGAACAAGTGGAACAGCTGTTAC-3’ |
| M1-MR | (FAM) 5’-ATCTGAAGGATTTGTACGTCTTGAATTACC-3’ |

Primer sequences for *P. falciparum* nested PCR targeting *msp1* K1

| Name | Sequence |
|---|---|

| | |
|---|---|
| M1-KF | Tail 5’-AAATGAAGAAGAAATTACTACAAAAGGTGC-3’ |
| M1-KR | (NED) 5’-GCTTGCATCAGCTGGAGGGCTTGCACCAGA-3’ |

Primer sequences for *P. falciparum* nested PCR targeting *msp2* FC27

| Name | Sequence |
|---|---|
| S tail – fw | 5’-GCTTATAATATGAGTATAAGGAGAA-3’ |
| M5 – rev | 5’- GCA TTG CCA GAA CTT GAA -3' FAM |

Primer sequences for *P. falciparum* nested PCR targeting *msp2* 3D7

| Name | Sequence |
|---|---|
| S tail – fw | 5’-GCTTATAATATGAGTATAAGGAGAA-3’ |
| N5 – rev | 5’- CTG AAG AGG TAC TGG TAG A -3' VIC |

Primer sequences for *P. falciparum* nested PCR targeting *glurp*

| Name | Sequence |
|---|---|
| GNF_fw | (PET) 5’-TGTTCACACTGAACAATTAGATTTAGATCA-3’ |
| G3_rev | TAIL 5’-TGTAGGTACCACGGGTTCTTG-3’ |

Primer sequences for *P. falciparum* PCR targeting microsatellites TA40, TA60, TA81 and PfPK2

| | Forward | Reverse |
|---|---|---|
| TA40 | 6FAM-TTTTGGTTTCCAAGGGATTG | gtgtcttTTAAGGCCACGAGGAAATTG |
| TA60 | HEX-CCAAGAGAAAGCGATCCTCA | gtgtcttTTTTTCCATCATATAAATTGGTATCT |
| TA81 | HEX-AGGGAAGGTGAGGAAAAGGA | gtgtcttTTCATACATTTCACACAACACAGG |
| PfPK2 | HEX-TCCTCAGACTGAAATGCATGA | gtgtcttCCTTTCATCGATACTACGATTATTTG |

Method 3: Targeted amplicon deep sequencing

Primer sequences for *P. falciparum* primary PCR targeting *ama1-D3*

| Name | Sequence |
|---|---|
| ama1-D3 (fw) | GTTTAATTAACAATTCATCATAC |
| ama1-D3 (rv) | GTGTTGTATGTGATGCTC |

Primer sequences for *P. falciparum* primary PCR targeting *cpmp*

| Name | Sequence |
|---|---|
| cpmp (fw) | CGATACAGGACATATAGA |
| cpmp (rv) | TTCAATAACATTTACTAGG |

Primer sequences for *P. falciparum* primary PCR targeting *csp*

| Name | Sequence |
|---|---|
| csp (fw) | ATCAAGGTAATGGACAAG |
| csp (rv) | ACTCAAACTAAGATGTGTTC |

Primer sequences for *P. falciparum* primary PCR targeting *cpp*

| Name | Sequence |
|---|---|
| cpp (fw) | TGTCTGAACCAAATTCAA |
| cpp (rv) | GAATTTGTCACATTTGATGA |

Primer sequences for *P. falciparum* primary PCR targeting *msp7*

| Name | Sequence |
|---|---|
| msp7 (fw) | GTATTATCAAAGGTAAAGGCA |
| msp7 (rv) | TTGCATAACTATAAACACCAT |

Primer sequences for *P. falciparum* nested PCR targeting *ama1-D3*

| Name | Sequence |
|---|---|
| ama1-D3_fw_linker | **GTGACCTATGAACTCAGGAGTC**TACTACTGCTTTGTCCCATC |
| ama1-D3_rv_linker | **CTGAGACTTGCACATCGCAGC**TCAGGATCTAACATTTCATC |

Primer sequences for *P. falciparum* nested PCR targeting *cpmp*

| | |
|---|---|
| cpmp_fw_linker | **GTGACCTATGAACTCAGGAGTC**CATAAGTCATTAAAATTTAT GGAT |
| cpmp_rv_linker | **CTGAGACTTGCACATCGCAGC**CGTTACTATCAAGATCGTTAATATC |

Primer sequences for *P. falciparum* nested PCR targeting *csp*

| Name | Sequence |
|---|---|
| csp_fw_linker | **GTGACCTATGAACTCAGGAGTC**AAATGACCCAAACCGAAATGT |
| csp_rv_linker | **CTGAGACTTGCACATCGCAGC**GGAACAAGAAGGATAATACCA |

Primer sequences for *P. falciparum* nested PCR targeting *cpp*

| Name | Sequence |
|---|---|
| cpp_fw_linker | **GTGACCTATGAACTCAGGAGTC**CAAGTTCACTTTTGGGAAATG |
| cpp_rv_linker | **CTGAGACTTGCACATCGCAGC**ATTACTACCTTTCAGCATATCCGA |

Primer sequences for *P. falciparum* nested PCR targeting *msp7*

| Name | Sequence |
|---|---|
| msp7_fw_linker | **GTGACCTATGAACTCAGGAGTC**ATGAACAAGAGATATCAACACA |
| msp7_rv_linker | **CTGAGACTTGCACATCGCAGC**TTAAATTGTTCATGGTATTCCTTA |

Primer sequences for *P. falciparum* adapter PCR

Forward:

AATGATACGGCGACCACCGAGATCTACACTCTTTCCCTACACGACGCTCTTCCGATCTXXXXXXXXGTGACCTAT GAACTCAGGAGTC

XXXXXXXX = forward barcode

GTGACCTATGAACTCAGGAGTC = linker forward

Reverse:

CAAGCAGAAGACGGCATACGAGATCGGTCTCGGCATTCCTGCTGAACCGCTCTTCCGATCTXXXXXXXXCTGA GACTTGCACATCGCAGC

XXXXXXXX = reverse barcode

CTGAGACTTGCACATCGCAGC = linker reverse

Forward barcode for *P. falciparum* adapter PCR forward

| Name | Sequence |
|---|---|
| Fw_2 | CTCTCTAT |
| Fw_3 | TATCCTCT |
| Fw_4 | AGAGTAGA |
| Fw_5 | GTAAGGAG |
| Fw_6 | ACTGCATA |
| Fw_7 | AAGGAGTA |

| | |
|---|---|
| Fw_8 | CTAAGCCT |
| Fw_9 | CCGAAGTA |
| Fw_10 | GAGCTGAA |
| Fw_11 | GCGAGTAA |

Primer sequences for *P. falciparum* adapter PCR reverse

| Name | Sequence |
|---|---|
| Rv_1 | TAAGGCGA |
| Rv_2 | CGTACTAG |
| Rv_3 | AGGCAGAA |
| Rv_4 | TCCTGAGC |
| Rv_5 | GGACTCCT |
| Rv_6 | TAGGCATG |
| Rv_7 | CTCTCTAC |
| Rv_8 | CAGAGAGG |
| Rv_9 | GCTACGCT |
| Rv_10 | CGAGGCTG |
| Rv_11 | AAGAGGCA |
| Rv_12 | GTAGAGGA |

Method 4: High-resolution melting analysis

Primer sequences for *P. falciparum* quantitative PCR-HRM targeting *msp1*

| Name | Sequence |
| --- | --- |
| *Msp1*HRM_F (fw) | TAGAAGATGCAGTATTGACAGGT |
| *Msp1*HRM_R (rv) | CAGCGTAAGATTTAGCATCTGAATC |

Primer sequences for *P. falciparum* quantitative PCR-HRM targeting *msp2*

| Name | Sequence |
| --- | --- |
| *Msp2*HRM_F (fw) | AGCAACACATTCATAAACAATGCT |
| *Msp2*HRM_R (rv) | TCCATGTTGTCCTGTACCTTTATTC |
